# RaptRanker: *in silico* RNA aptamer selection from HT-SELEX experiment based on local sequence and structure information

**DOI:** 10.1101/2019.12.31.890392

**Authors:** Ryoga Ishida, Tatsuo Adachi, Aya Yokota, Hidehito Yoshihara, Kazuteru Aoki, Yoshikazu Nakamura, Michiaki Hamada

## Abstract

Aptamers are short single-stranded RNA/DNA molecules that bind to specific target molecules. Aptamers with high binding-affinity and target specificity are identified using an *in vitro* procedure called high throughput systematic evolution of ligands by exponential enrichment (HT-SELEX). However, the development of aptamer affinity reagents takes a considerable amount of time and is costly because HT-SELEX produces a large dataset of candidate sequences, some of which have insufficient binding-affinity. Here, we present RNA aptamer Ranker (RaptRanker), a novel in *silico* method for identifying high binding-affinity aptamers from HT-SELEX data by scoring and ranking. RaptRanker analyzes HT-SELEX data by evaluating the nucleotide sequence and secondary structure simultaneously, and by ranking according to scores reflecting local structure and sequence frequencies. To evaluate the performance of RaptRanker, we performed two new HT-SELEX experiments, and evaluated binding affinities of a part of sequences that include aptamers with low binding-affinity. In both datasets, the performance of RaptRanker was superior to Frequency, Enrichment and MPBind. We also confirmed that the consideration of secondary structures is effective in HT-SELEX data analysis, and that RaptRanker successfully predicted the essential subsequence motifs in each identified sequence.

## 1 Introduction

Aptamers are chemically-synthesized short single-stranded RNA/DNA molecules that can bind various targets with high binding affinity and specificity. Aptamers have mainly two characteristics that distinguish them from other nucleic acid drugs. First, aptamers recognize the tertiary structure of target molecules, while other oligonucleotide drugs target mRNA molecules through their primary structures only. Second, aptamers can target extracellular molecules as well as intracellular molecules. Unlike other nucleic acid drugs, aptamers can be generated against various types of target molecules such as transcription factors [1, 2, 3, 4], proteins and their complexes [5], small organic molecules [6, 7], viruses [8], and cells [9, 10, 11]. Moreover, aptamer production by chemical synthesis offers easy manufacturing, batch-to-batch reproducibility, and long shelf life. Considering these features, aptamers show potential as replacements of antibodies in analytical, diagnostic, and therapeutic applications.

Aptamers bind their targets by forming tertiary structures that provide spatial complementarity [12]. Therefore, it is important to consider the structural features (e.g., RNA secondary structures) of aptamers when predicting their binding properties. The information regarding the secondary structure is utilized in aptamer development strategies such as motif definition and strategic truncation [13], *in silico* screening to reduce initial pool size [14], and searching the sequence space for potent aptamers [15, 16]. Although a variety of software is available for predicting secondary structures of aptamers (e.g., [17, 18]), only few analyze secondary structure information for the calculation of binding potentials of RNA aptamers.

Recently, aptamers with high binding-affinities were identified using HT-SELEX method [19, 1, 20]. HT-SELEX is an in *vitro* experimental method that improves the conventional SELEX method [21, 22] by including next generation sequencing (NGS) technology so that a larger amount of nucleotide sequence information can be processed. In (HT-)SELEX procedures, the aptamers with high binding-affinities are enriched by repeating the “Amplify only sequences that bind target” procedure, starting with large amounts of random RNA/DNA sequences. First, an RNA/DNA pool with 10^12^ to 10^15^ random nucleotide sequences called round 0 pool (0R pool) or initial pool is generated. Then, high binding-affinity aptamers are identified by performing repeated rounds of a five-step procedure as shown in figure 1. In HT-SELEX method, aptamers obtained in not only the final round but also in some of the previous rounds including the 0R pool are sequenced. Since a vast number of sequences are generated by HT-SELEX [23, 24, 25], an efficient candidate selection method is required to reduce the number of non-binding clones, decrease experiment time, and minimize material cost.

**Figure 1:**
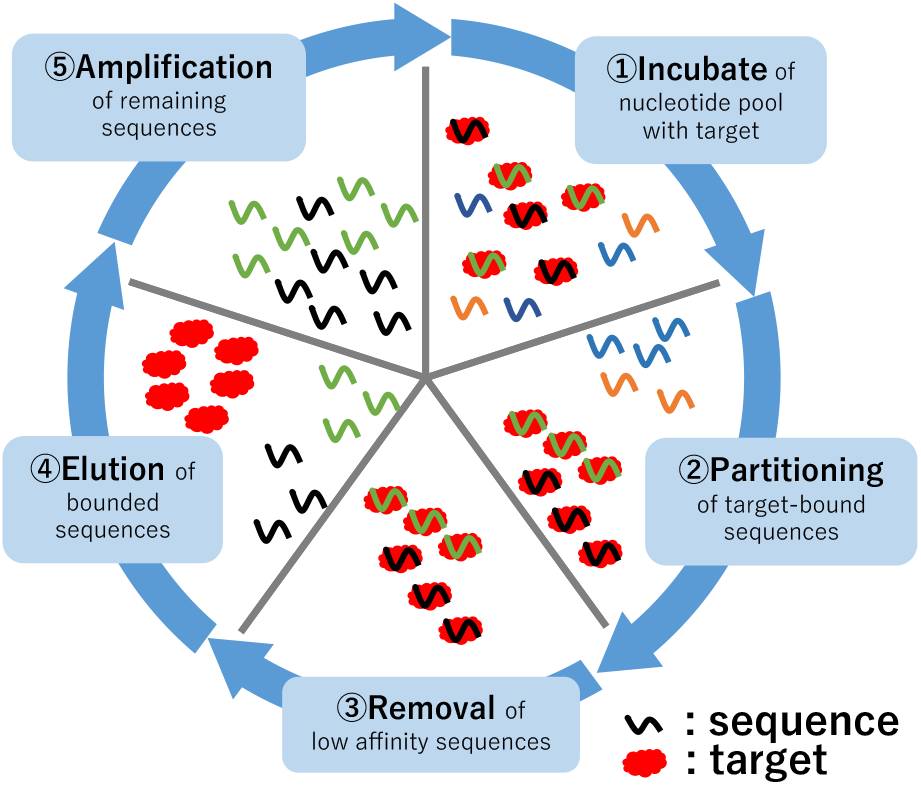
Diagram of HT-SELEX method. HT-SELEX method identifies high binding-affinity aptamers by performing repeated rounds of a five-step procedure. 1. Addition of the target molecule to the RNA/DNA pool and incubation. 2. Partitioning of the target molecule-RNA/DNA complex and unbound RNA/DNA. 3. Removal of unbound RNA/DNA. 4. Elusion of target-bound RNA/DNA. 5. Amplification of selected RNA/DNA.

There have been some previous *in silico* studies to identify the high binding-affinity sequences from HT-SELEX data by various analyses. Frequency, Enrichment [26], and MPBind [27] were developed to achieve this purpose by scoring and ranking the candidate sequences. Frequency calculates the frequency of appearance for each sequence. Enrichment calculates transition of frequency between consecutive rounds for each candidate sequence. MPBind performs statistical analysis based on changes in relative frequencies of all k-mer units. However, none of them includes aptamer structure data in their analyses. In contrast, AptaTRACE [13] traces the transition of secondary structure between HT-SELEX rounds for each k-mer and predicts the binding motif independent of frequency data. However, AptaTRACE does not provide a ranking for the candidates. APTANI^2^ [28] ranks candidates and identifies relevant structural motifs through the calculation of a score that considers secondary structures of aptamers. However, the score focuses on structural stability rather than structural similarity. In addition, APTANI^2^ uses only one round of HT-SELEX as its input.

Another challenge in software development is that there are only a few public HT-SELEX datasets. In aptamer-related research, only the aptamers that show high binding-affinity are reported; research reports rarely disclose the complete (HT-)SELEX data generated throughout the selection process. In many cases, even when the sequence data is made available, only the data from the final round is shared and the data generated in intermediate rounds of HT-SELEX procedure is not available. It should also be noted that there is little information about the candidate sequences that turn out to lack binding activity. This data of “aptamers with low binding-affinity” is highly valuable for evaluating the performances of *in silico* HT-SELEX analysis methods.

Here we present RNA aptamer Ranker (RaptRanker), a novel in *silico* method for identifying high binding-affinity aptamers from HT-SELEX data by scoring and ranking. RaptRanker determines unique sequences from all HT-SELEX rounds and clusters all subsequences of unique sequences by similarity based on both nucleotide sequence and secondary structure features. Then, RaptRanker identifies high binding-affinity aptamers by calculating the average motif enrichment (AME), which is a score assigned to each unique sequence based on frequency of subsequence clusters. We performed two new HT-SELEX experiments and evaluated sequence sets that include aptamers with low binding-affinity using surface plasmon resonance (SPR) assay. In both HT-SELEX dataset analyses, performance of RaptRanker was superior to Frequency, Enrichment, and MPBind.

## 2 MATERIALS AND METHODS

RaptRanker clusters the subsequences of unique sequences based on similarity in both nucleotide sequence and secondary structure. Then, RaptRanker assigns a score for each subsequence and each unique sequence using the cluster data. Finally, RaptRanker ranks all unique sequences by score (Figure 2). Importantly, the score calculated by RaptRanker reflects particular characteristics of aptamers and SELEX experiment, such as nucleotide sequence similarity, secondary structure similarity, and frequency information.

**Figure 2:**
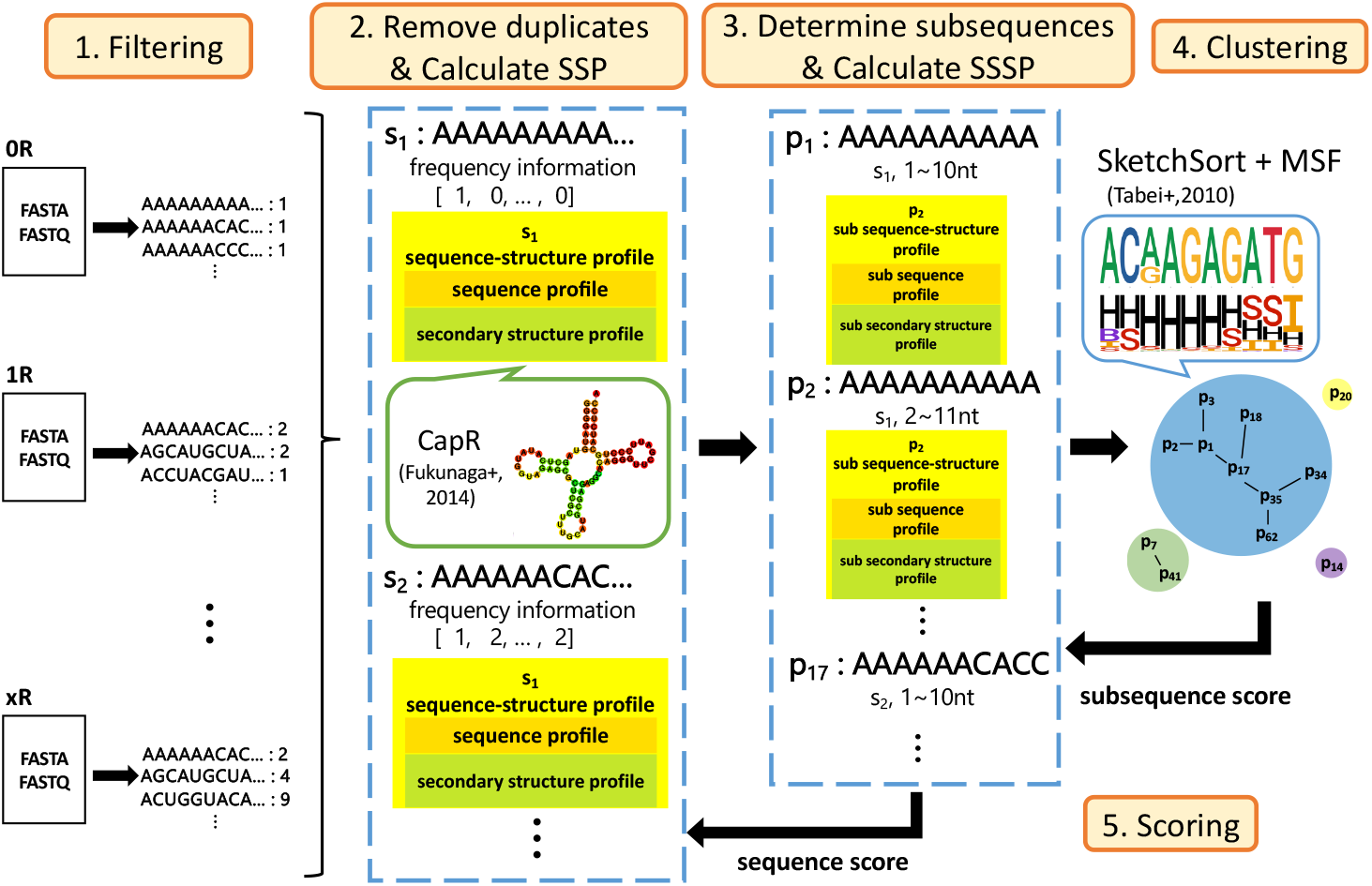
An illustration of RaptRanker algorithm. First, RaptRanker determines the sequences to be analyzed from all the input FASTA/FASTQ files of HT-SELEX experiments by filtering. Second, RaptRanker extracts unique sequences by removing duplicates, and computes sequence-structure profiles (SSPs) which represents both sequence and structure information. Third, RaptRanker determines subsequences of unique sequences, and constructs subsequence-structure profile (SSSP) by decomposing SSPs. Forth, RaptRanker clusters subsequences (from all the rounds) by constructing Minimum Spanning Forest (MSF), based on similar pairs of SSSPs calculated by SketchSort. Finally, sequence scores are computed based on clustering. Each cluster obtained by RaptRanker can be regarded as a motif because it is a collection of similar subsequences of nucleotide sequence and secondary structure.

Specifically, RaptRanker identifies high binding-affinity aptamers through the five steps shown in figure 2: (i) import of the sequence data from FASTA or FASTQ input file, (ii) determination of unique sequences by removing duplicates and computation of sequence-structure profiles (SSPs) representing nucleotide sequence and secondary structure features as predicted by CapR [17], (iii) derivation of subsequences sequentially from all unique sequences, and generation of sub-sequence-structure profiles (SSSPs) representing nucleotide sequence and secondary structure features for each subsequence, (iv) enumeration of all subsequence pairs matched based on similarity in sub-sequence-structure profiles using SketchSort [29] and clustering of subsequences by constructing minimum spanning forest (MSF) from all similar subsequence pairs, and (v) calculation of the scores and ranking unique sequences.

The details of each step are described in the following sections. (See Table S1 for notations utilized below.)

### 2.1 Filtering

RaptRanker determines sequences to be analyzed by two-step filtering. First, only the sequences whose primer-binding region is the same as the design are extracted. Then, the random regions whose length is within the user-defined limits are extracted. The sequences (i.e., the set of random regions) that pass the filtering criteria are subjected to further analyses.

In the first filtering, the sequences from FASTA or FASTQ files are imported and only the sequences whose primer binding regions exactly matches the design are extracted. Both ends of sequences obtained from the SELEX experiment include fixed nucleotide sequences necessary for amplification by PCR method. These sequences are called forward primer binding region (5’ end) and reverse primer binding region (3’ end). RaptRanker extracts only the sequences whose both forward and reverse primer binding regions exactly match the user-defined parameters *forward_primer* and *reverse_primer*.

In the second filtering, the random regions whose lengths are between the upper and lower limits are extracted. RaptRanker uses the user-defined parameters *minimum_length* and *maximum_length*, and extracts only the random regions whose lengths *l* are *minimum_length* ≤ *l* ≤ *maximum_length*.

### 2.2 Removal of duplicates and computation of sequence-structure profiles (SSPs)

#### 2.2.1 Removal of duplicates

RaptRanker determines the unique sequences by removing duplicates present in the imported sequence dataset in advance to reduce the calculation cost. First, for each round, count the appearance of each sequence and remove duplicates with radix sort [30]. After that, summarize the results of each round, and remove duplicates with radix sort again. This result is the unique sequences from all inputted HT-SELEX data, and each unique sequence has the frequency information on each round.

In the following, let *S* be the set of all unique sequences, and let *s* be a unique sequence in *S*.

#### 2.2.2 Computation of sequence-structure profiles (SSPs)

For each unique sequence *s*(∈ *S*), RaptRanker computes sequence-structure profile (SSP) that represents nucleotide sequence and secondary structure features as predicted by CapR [17]. SSP consists of a sequence profile (4 × unique sequence length) representing the nucleotide sequence and a secondary structure profile (6 × unique sequence length) representing the secondary structure (Figure 3a).

**Figure 3:**
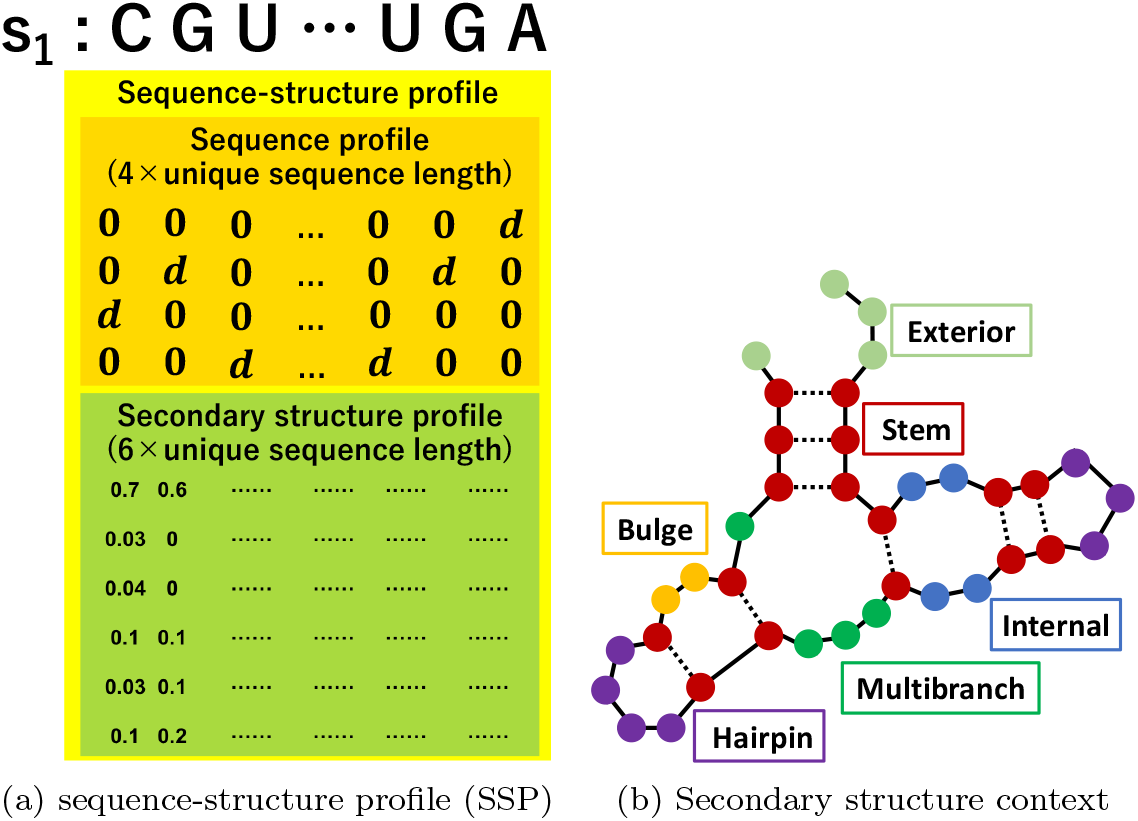
(**a**) Diagram of sequence-structure profile (SSP). SSP is computed for each unique sequence, and consists of a sequence profile (4 × unique sequence length) representing the nucleotide sequence and a secondary structure profile (6 × unique sequence length) representing the secondary structure. The sequence profile is a matrix computed by converting the nucleotide sequence of each unique sequence into a one-hot vector using a real value *d* ∈ ℝ. The secondary structure profile is a matrix computed by converting the data obtained from CapR. (**b**) An example image of RNA secondary structure. The dashed lines represent base pairs. The secondary structure features predicted by CapR are classified into six types (Bulge, Exterior, Hairpin, Internal, Multibranch, Stem).

The secondary structure is predicted for each unique sequence that includes a part of primer binding regions in addition to the random region. In this way, CapR can include the effect of the primer binding region on the secondary structure in the analysis. RaptRanker uses user-defined parameters *add-forward-primer* and *add_reverse_primer* in addition to and separate from parameters for filtering (Supplementary Tables S2 and S3).

Sequence profile is a (4 × unique sequence length) matrix representing the nucleotide sequence information of each unique sequence *s*(∈ *S*). The sequence profile is computed by converting the nucleotide sequence data into a one-hot vector using a real value d ∈ ℝ as *A* = [*d*, 0,0,0]^*T*^, *G* = [0,*d*,0,0]^*T*^, *C* = [0,0,*d*,0]^*T*^, *U* = [0,0,0,*d*]^*T*^ (Figure 3a). The real value *d* is a user-defined parameter *weight* representing the weight of the sequence information in subsequent clustering. If the value of *d* is large, the sequence information becomes more emphasized than the secondary structure information, and therefore more strict matching of the nucleotide sequences between entries is required to be clustered together.

Secondary structure profile is a (6 × unique sequence length) matrix representing the secondary structure information of each unique sequence *s*(∈ *S*). Secondary structure profile is computed by converting the data obtained from CapR [17] into a matrix (Figure 3a). CapR is a software which calculates the probability of each base of RNA sequence being a part of a specific type of secondary structure. Secondary structure features predicted by CapR are classified into six types (Bulge, Exterior, Hairpin, Internal, Multibranch, Stem) (Figure 3b). Since the output of CapR is a probability value, the sum of six rows in each column of the secondary structure profile is 1.0.

### 2.3 Determination of subsequences and computation of sub-sequence-structure profiles (SSSPs)

#### 2.3.1 Determination of subsequences

RaptRanker determines the list of subsequences. Subsequences are determined by cutting out a fragment from a unique sequence *s*(∈ *S*), and repeating this process, each time shifting the location of the fragment by one base. Each subsequence is indexed individually, and contains the index information of unique sequence it was obtained from. Each subsequence also contains its original position on the corresponding sequence. Even if the nucleotide sequences are identical, they are distinguished by this information and treated separately. RaptRanker includes a user-defined parameter *wide* as window length.

In the following, let *P* be the set of all subsequences, and let *p* be a subsequence. In addition, let *D_s_* be the set of subsequences generated from unique sequence *s*. In this case, ⋃_*s*∈S_ *D_s_* = *P* holds. Also, since each subsequence *p*(∈ *P*) is treated separately, 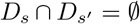 holds for any set of two unique sequences *s*, *s′*(∈ *S*).

#### 2.3.2 Computation of sub-sequence-structure profiles (SSSPs)

For each subsequence *p*(∈ *P*), RaptRanker computes sub-sequence-structure profile (SSSP) representing nucleotide sequence and secondary structure features of the subsequence by dividing SSPs into sub-profiles of a fixed length (Figure 4).

**Figure 4:**
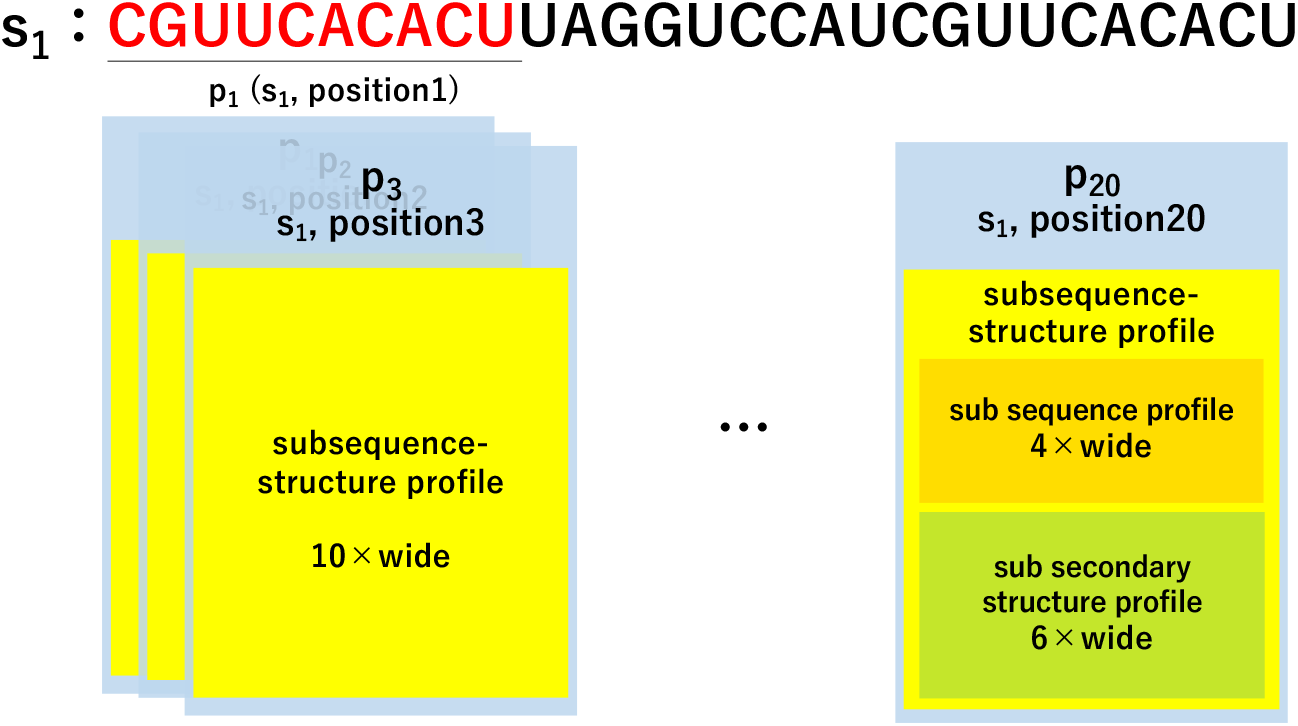
Diagram of determination of sub-sequence-structure profile (SSSP) in case of *wide* = 10. Each subsequence *p*(∈ *P*) is indexed individually and contains the index information of the unique sequence it was obtained from, its original position on the corresponding sequence, and SSSP. In this example, the nucleotide sequence of *p*_1_ and *p*_20_ are identical (“CGUUCACACU”), but they are treated separately.

### 2.4 Clustering subsequences

#### 2.4.1 Enumeration of all similar subsequence pairs

RaptRanker enumerates all similar subsequence pairs based on SSSP similarity using SketchSort [29]. SketchSort is a software that can enumerate similar pairs in a large number of high dimensional vectors rapidly and approximately. RaptRanker converts each SSSP into a row vector to be provided as an input to SketchSort. SketchSort requires two parameters; an upper limit of cosine distance between vectors *cosdist*, and an upper limit of the expectation value for false negatives *missing_ratio.*

#### 2.4.2 Clustering of all subsequences

RaptRanker clusters all subsequences by constructing a MSF from all similar subsequence pairs. MSF can be calculated efficiently by Kruskal’s algorithm [31] and UnionFind (Appendix B).

Since each MST from MSF becomes a cluster, clusters do not overlap each other in, i.e. a subsequence does not belong to more than one cluster.

In the following, let 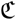 be the set of clusters, and let *C* be a cluster. In this case, since each cluster *C* is a set of subsequences and *P* is the set of all subsequences, 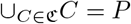 holds. Also, since clusters do not overlap each other, 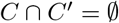 holds for any set of two clusters 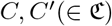.

### 2.5 Calculation of score

RaptRanker calculates average motif enrichment (AME), a score for predicting binding affinity, for each unique sequence *s*(∈ *S*) based on the clustering results through three steps. (i) Calculation of subsequence frequency for each subsequence *p*(∈ *P*). (ii) Calculation of motif frequency and motif enrichment for each cluster 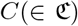, and define motif enrichment of each cluster 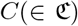 as the score of each subsequence in that cluster *p*(∈ *C*). (iii) Calculation of AME for each unique sequence *s*(∈ *S*) based on motif enrichment of subsequences *p*(∈ *D_s_*).

#### 2.5.1 Calculation of subsequence frequency

Subsequence frequency is calculated for each subsequence *p*(∈ *P*) and for each round. It is calculated by dividing the number of occurrences of each subsequence by the total number of occurrences of subsequences. Subsequence frequency of a subsequence *p* at round *x* is

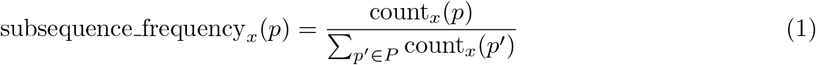

where count_*x*_ (*p*) is the occurrence of the subsequence *p* at round *x*.

#### 2.5.2 Calculation of motif frequency and motif enrichment

Motif frequency is calculated for each cluster 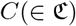 and for each round. Motif frequency is the sum of the subsequence frequencies of subsequences in a cluster. Motif frequency of a cluster *C* at round *x* is

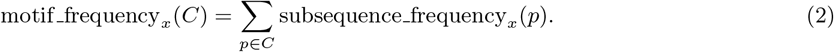

Then, RaptRanker calculates motif enrichment for each cluster 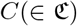) and for each round. Motif enrichment is calculated for a specific round by dividing the motif frequency in that round by the motif frequency in the previous round. Motif enrichment of a cluster *C* at round *x* is

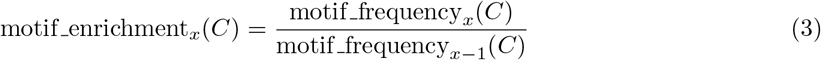

for motif_frequency_*x*−1_(*C*) ≠ 0 and round *x* is not the first round; the score is undefined otherwise.

RaptRanker defines motif enrichment of each cluster 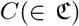 as the score of each subsequence in that cluster *p*(∈ *C*).

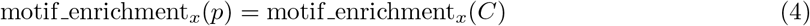

where *p* ∈ *C*.

#### 2.5.3 Calculation of Average Motif Enrichment (AME)

AME is calculated for each unique sequence *s*(∈ *S*) and for each round. AME is the average of the motif enrichment values of subsequences from a unique sequence. The AME of a unique sequence s at round *x* is defined as

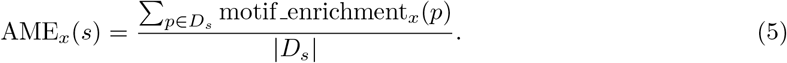

The score is undefined for the first round.

### 2.6 Newly acquired experimental data in this study

Because the number of existing datasets is not sufficient for evaluating our method, we performed two new HT-SELEX experiments, and selected evaluation sequence sets that include aptamers with low binding-affinity as determined by surface plasmon resonance (SPR) assay. The target molecules of HT-SELEX experiments were human recombinant transglutaminase 2 (TG2; Cat. No. 4376-TG) and human recombinant integrin alpha V beta 3 (*α*V*β*3; Cat. No. 3050-AV). In the following, we refer the data from TG2 experiment as Data1, and the data from *α*V*β*3 experiment as Data2. The sequence data are available on DRA009383 and DRA009384, and evaluation of sequence sets in each dataset are shown in Supplementary Tables S4 and S5.

TG2 and *α*V*β*3 were purchased from R&D systems. Z-Gln-Gly (Cat. No. C6154) and Hydroxylamine hydrochloride (Cat. No. 169417) were obtained from Sigma-Aldrich. All other buffers and chemicals were of special grade.

#### 2.6.1 HT-SELEX

Selection of aptamers was followed by a previously-reported SELEX method [32] with some modifications. The sequence of the first template for Data1 and Data2 were 5’ -- TCACACTAGC ACGCATAGG --30N-- CATCTGACCT CTCTCCTGCT CCC --3’ and 5’-- GAGGATCCAT GTATGCGCAC ATA --40N-- CTTCTGGTCG AAGTTCTCCC --3’, respectively. The following primers were used. Data1-Forward, 5’-- TAATACGACTC ACTATAGGGAG CAGGAGAGAG GTCAGATG --3’; Data1-Reverse, 5’-- TCACACTAGC ACGCATAGG --3’; Data2-Forward, 5’-- CGGAATTCT AATACGACT CACTATAGGG AGAACTTCG ACCAGAA --3’; Data2-Reverse, 5’-- GAGGATCCAT GTATGCGCA CATA --3’. The oligonucleotide libraries containing 2’-fluoro-pyrimidine nucleotides were prepared by transcription using a mutant T7 RNA polymerase. The binding buffer used for selection contained 145 mM NaCl, 5.4 mM KCl, 5 mM MgCl2, 0.05% Tween20, and 20 mM Tris-HCl (pH 7.6). For Data2 SELEX, 1 mM MnCl2 was used instead of MgCl2. The recombinant proteins were immobilized on NHS-activated Sepharose (17-0906-01, GE Healthcare) and bound nucleotides were released with 6 M Urea. After selection, nucleotide pools were sequenced using an Ion PGM Hi-Q View Sequencing Kit (A30044, Thermo Fisher Scientific).

#### 2.6.2 Surface plasmon resonance (SPR) assay

The SPR assays were performed essentially as described previously using the Biacore T200 instrument (GE Healthcare) [32]. Aptamers were prepared in 16-merpoly(A)-tailed form by in vitro transcription. A 5’-biotinylated dT16 oligomer was bound to the surface of the streptavidin sensor chip (BR100531, GE Healthcare) of active and reference flow cells. The poly(A)-tailed RNA was immobilized to the active flow cell by complementary hybridization to the dT16 oligomer. The recombinant protein in the binding buffer was injected to both flow cells of the sensor chip. Data was obtained by subtracting the reference flow cell data from the active flow cell data. Maximum response after injection was used for determination of true binding clone. Experimental parameters, including material concentrations, flow rate, contact time, and criteria of true binding clone were optimized for each target (see Table 1). To regenerate the sensor chip, bound material was completely removed by injecting 6 M urea. The detailed results are shown in Supplementary Figures S1a–S2c.

**Table 1:** Parameters of surface plasmon resonance (SPR) assay

| Dataset | Aptamer<br>(nM) | Recombinant<br>protein (nM) | Flow rate<br>(μL/min) | Contact<br>time (s) | Criteria for<br>True (RU) |
| --- | --- | --- | --- | --- | --- |
| Data1 | 300 | 300 | 20 | 60 | > 25 |
| Data2 | 100 | 100 | 30 | 60 | > 25 |

#### 2.6.3 Transglutaminase assay

Inhibitory activities of aptamers in Data1 was assessed by following procedures. Forty microliter of reaction mixture comprising 200 ng TG2, 50 mM Z-Gln-Gly, 100 mM hydroxylamine, 10 mM DTT, 10 mM CaCl2, and different concentrations of aptamer (0-1000 nM) in SELEX buffer was used in the assay. After incubation at 37 °C for 3 hours, 160 *μ*L of assay reagent (370 mM FeCl3, 200 mM trichloroacetate, 670 mM HCl) was added and the solution was centrifuged for 2 min at 20,000 ×*g*. The supernatant was transferred to a microplate, and the resulting color was measured at 525 nm. IC50 value was calculated by XLfit.

## 3 Results and Discussion

We used data from rounds 0 to 4 on Data1, and 3 to 6 on Data2 as input for analysis. Data1 comprises data from rounds 0 to 8, but we did not include data from 5 to 8 in the analysis because we confirmed that the sequences evaluated as False were significantly amplified those rounds (see the Supplementary material, Section 3.4.3, for the details).

In the following, we used the parameters (*wide*=10, *weight*=0.5, *cosdist*=0.001, *missing_ratio*=0.00001) on RaptRanker for both data. Other data-specific parameters (related to filtering) are listed in supplementary tables S2 and S3.

### 3.1 Comparison of performances for the identification of high binding-affinity aptamer

To assess the performance of high binding-affinity aptamer identification, we compared “True Positive Rate (TPR) when False Positive Rate (FPR) is 0” metrics of two HT-SELEX datasets. This evaluation metrics corresponds to the rise of the ROC curve, and it considers the practical use that the binding-affinity is experimentally confirmed only for high scoring sequences (details are on section 3.4.2). Among the existing methods, Frequency and Enrichment were re-implemented and calculated by RaptRanker (Appendix D).

We compared each approach in the final round of both datasets. RaptRanker showed the highest TPR in both datasets. So, RaptRanker identified the high binding-affinity aptamers with the highest performance among all three methods tested (Table 2). Here, the result of MPBind in Data1 was 0.0. MPBind is a method that can use control-round (SELEX round without adding target molecules) information. However, in this case, control-round information is not available in both data sets. This may be have caused the low performance of MPBind.

**Table 2:** Result of the performance comparison for the identification of high binding-affinity aptamers

| Methods | Data1.4R TPR | Data2.6R TPR |
| --- | --- | --- |
| RaptRanker | **0.524** | **0.361** |
| Enrichment | 0.238 | 0.333 |
| Frequency | 0.143 | 0.194 |
| MPBind | 0.000 | 0.250 |

We compared the performance of high binding-affinity aptamers identifying on two HT-SELEX datasets. The evaluation metrics is “True Positive Rate (TPR) when False Positive Rate (FPR) is 0”. We compared at 4R and 6R, the last round of inputs for each dataset. For each dataset, the highest values are shown in bold.

### 3.2 Validation of the effectiveness of analyzing both sequence and secondary structure

We validated the effectiveness of considering both nucleotide sequence and secondary structure. We compared binding-affinity prediction performances in a manner similar to that explained in the previous section. We included three types of analyses; first one is the analysis based on sequence similarity only, second one on secondary structure similarity only, and the last one on both parameters.

For analyses relying on sequence similarity only, we calculated Average K-mer Enrichment (AKE) in a similar way we calculated Average Motif Enrichment (AME) based on RaptRanker’s clustering (Appendix H); AKE is calculated based on k-mer, which is a set of subsequences with matching nucleotide sequences, while AME is calculated based on cluster RaptRanker clusters. Here, we calculated AKE for 10-mer because we set *wide* = 10 in RaptRanker.

As a method considering only secondary structure similarity, we calculated AME with *weight* = 0.0. By setting the parameter *weight* = 0.0, RaptRanker clusters the subsequences based on secondary structures similarity only. The AME calculated based on this clustering can be regarded as a score that predicts binding-affinity using secondary structure similarity only.

We evaluated the performances of each approach using the data from the final rounds of both datasets. Data generated by the method of analyzing similarities in both sequence and structural features showed the highest TPR in both datasets (Table 3). In Data1, the analysis based on “*structure* similarity only” and the analysis based on “both similarities” showed the same TPR values. In contrast, in Data2, the analysis based on “*sequence* similarity only” and the analysis based on both similarities showed the same TPR values. These results suggest that there are several types of HT-SELEX data, and that both sequence similarity and structure similarity should be considered in HT-SELEX data analysis. These results also suggest that RaptRanker is particularly useful in cases where the existing methods based on sequence similarity only cannot provide sufficient identification performance.

**Table 3:**
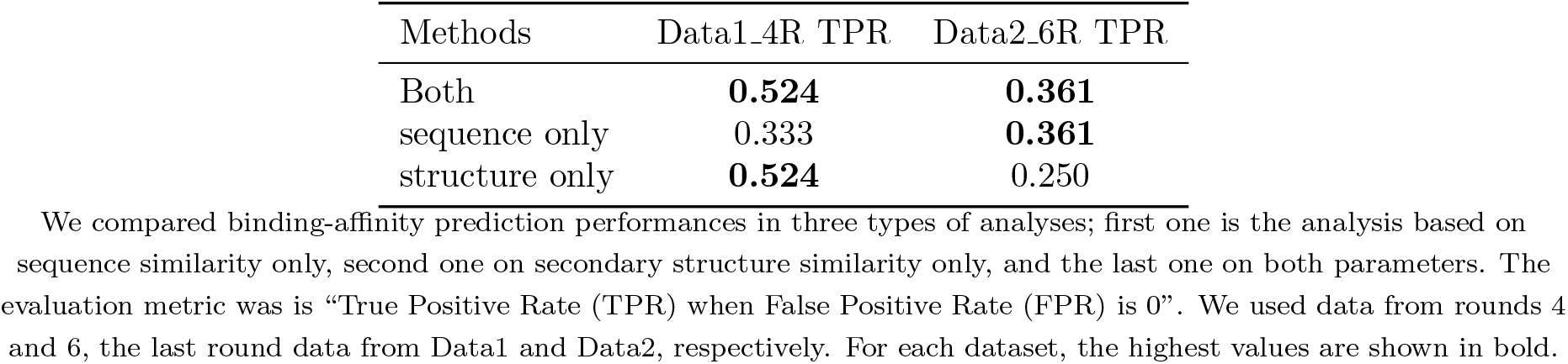
Result of validation of the effectiveness

### 3.3 Prediction of binding motifs in identified sequences

We acquired the truncated sequences from both datasets, and compared the truncated sequences with the binding motifs predicted by RaptRanker.

#### 3.3.1 Truncation of sequences

For both Data1 and Data2, we generated several truncated forms of candidate aptamers according to Transglutaminase assay and SPR assay (Supplementary Figures S3 and S4). Minimal active sequences were defined as 25 nt and 36 nt (Table 4). In addition, for Data2, we synthesized a series of single nucleotide deletion mutants of aptamers to predict binding motifs (Supplementary Figure S5). As a result, we estimated that “UACGU---CUG” is a binding motif.

**Table 4:**
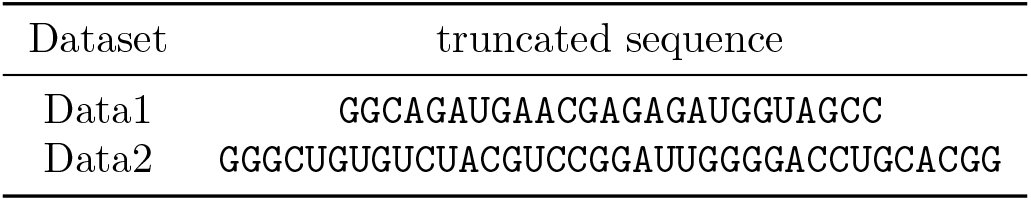
Truncated sequences

#### 3.3.2 Binding motif prediction with RaptRanker

The highest scoring subsequence from the sequence identified by RaptRanker can be regarded as the binding motif. Since RaptRanker calculates motif_enrichment_*x*_(*p*) for each subsequence *p*(∈ *D_s_*) from each unique sequence *s*(∈ *S*), the cluster *C* to which the highest scoring subsequence belongs *p* (the cluster *C* holds *C* ∋ *p* where the *p* holds max_*p*∈*D_s_*_ motif_enrichment_*x*_(*p*)) can be regarded as the binding motif of the unique sequence s. Here, we selected the unique sequences with true binding-affinity (Table 5).

**Table 5:**
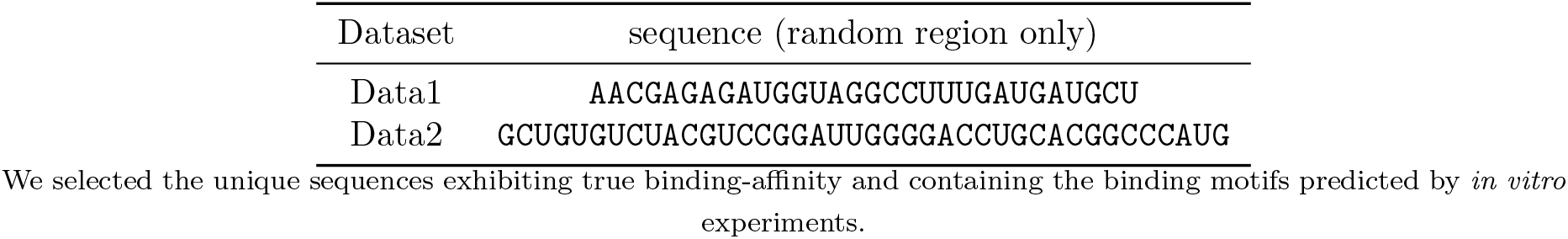
The unique sequences for binding motif prediction

#### 3.3.3 Comparison of motifs predicted by RaptRanker with experimentally-determined truncated sequences

We compared the truncated sequences and the binding motifs predicted by RaptRanker. In both datasets, the binding motifs predicted by RaptRanker were found to be parts of the truncated sequences (Figure 5ab). This result suggests that RaptRanker can predict the essential subsequence in each identified sequence. This information can be used for the optimization of identified sequences.

**Figure 5:**
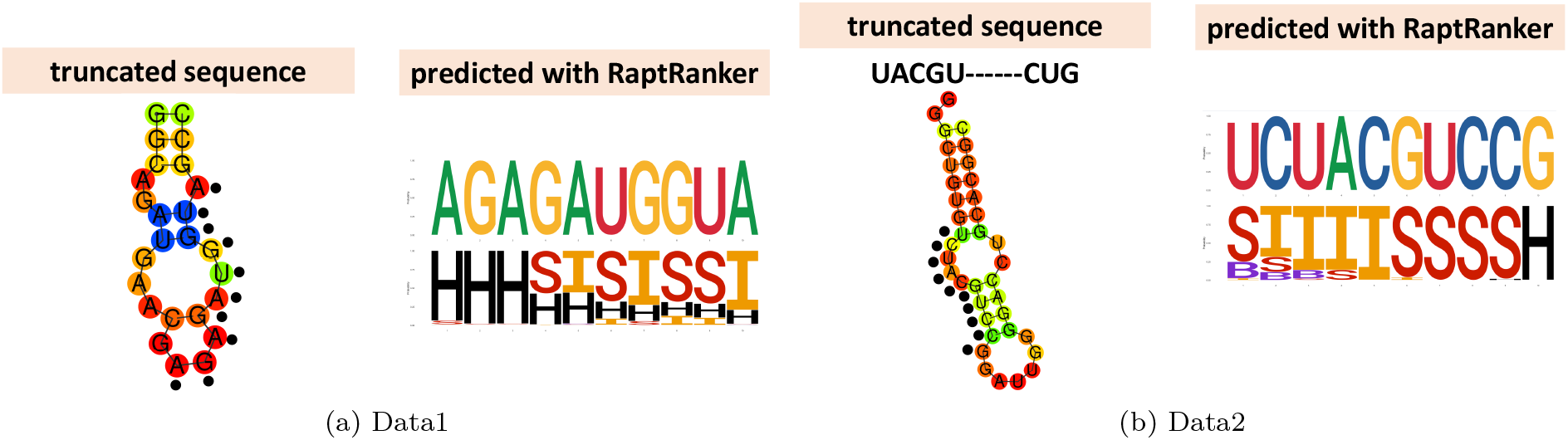
We compared the truncated sequences and the binding motifs predicted by RaptRanker. Black dots in truncated sequences show the nucleotide sequences that are identical to binding motifs predicted with RaptRanker. H, S, I, and B indicate hairpin, stem, internal loop, and bulge loop, respectively (cf. Figure 3b).

In addition, for Data2, we compared experimentally-determined motifs to the motifs predicted by RaptRanker, and found them to be similar (Figure 5b). However, short and discontinuous motifs such as those shown in figure 5b may affect the binding-affinity prediction performance of RaptRanker negatively because such discontinuous motifs are recognized as two different motifs in RaptRanker. Moreover, short motifs are harder to detect when the actual motif length is shorter than the value of *wide* in RaptRanker. Considering these shortcomings, it is conceivable that the binding affinity prediction performance of RaptRanker in Data2 did not improve as much as in Data1 due to presence of discontinuous and short motifs.

### 3.4 Further discussion

#### 3.4.1 Significance of clustering all subsequences simultaneously

RaptRanker enables the computation of motif enrichment by clustering all subsequences obtained in all rounds simultaneously. That is, RaptRanker takes advantage of data generated in multiple rounds of HT-SELEX by clustering all subsequences at once.

In the case of clustering all subsequences at once, clustering is not affected differences among rounds even if “Subsequences from sequences not observed in some rounds” are present. RaptRanker determines all unique sequences *S* from all rounds, and clusters all their subsequences *P*. In this case, those subsequences only affect the calculation of motif frequency and motif enrichment (Figure 6a).

**Figure 6:**
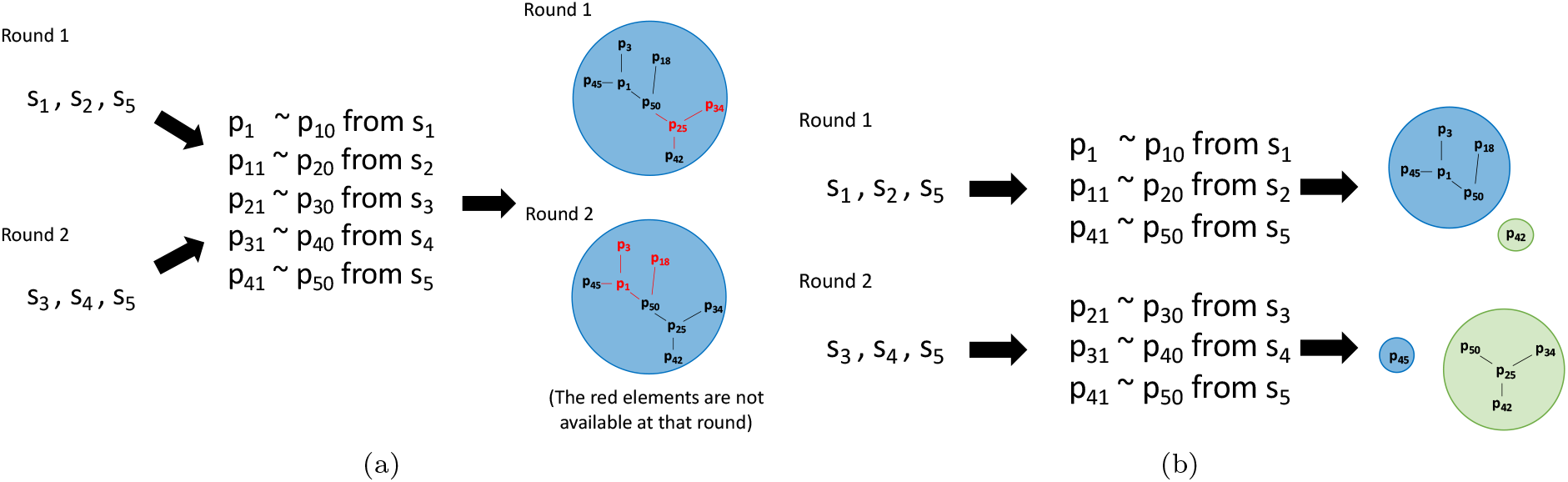
**(a)** In the case of clustering subsequences from all rounds simultaneously, the clustering is not affected by differences among rounds even if “Subsequences from sequences not observed in some rounds” are present. **(b)** In the case of clustering subsequences on each round, it is necessary to follow individual clusters across rounds. In this case, cluster fragmentation or integration may occur between rounds.

In contrast, in the case of clustering subsequences on each round, it is necessary to follow individual clusters through rounds. Cluster fragmentation or integration may occur between rounds in some cases. Such events make cluster mapping and computation of motif enrichment difficult (Figure 6b).

Since RaptRanker clusters all subsequences from all rounds simultaneously, it can track the transition of motif frequency and compute motif enrichment. As an example, we visualized the transition of motif frequency for the motifs shown in figure 5ab (Figure 7). The motif on Data1 is not observed in rounds 0 to 2, but it was highly enriched between rounds 3 and 4 (Figure 7a). In contrast, the motif in Data2 was enriched smoothly between rounds 3 and 6 (Figure 7b).

**Figure 7:**
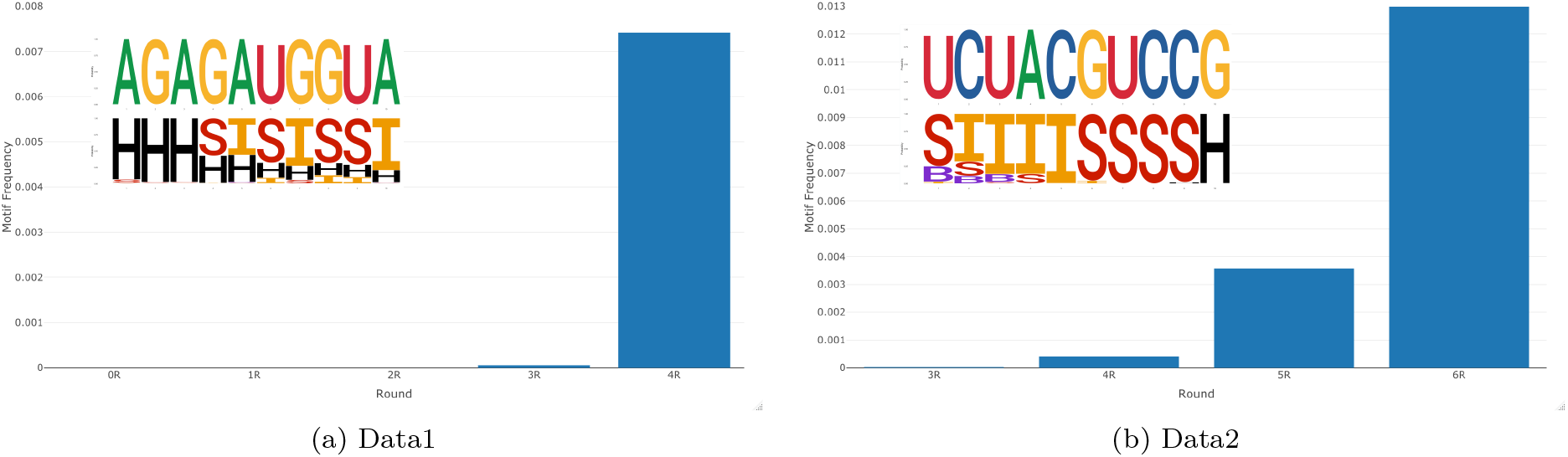
Transition of motif frequency across rounds. In each subfigure, horizontal axis and vertical axis show round number and motif frequencies, respectively. The motifs are predicted by RaptRanker. RaptRanker can track the transition of motif frequency by clustering subsequences from all rounds simultaneously.

#### 3.4.2 Evaluation metrics of binding affinity prediction performance

We used TPR when FPR is 0 as the metric for evaluation of binding-affinity prediction performance. This metric is based on the approach of ROC enrichment [33]. ROC enrichment is used for evaluation of protein-ligand virtual screening and calculated by dividing TPR by FPR. According to Jain and Nicholls, ROC enrichment should be reported in four points of FPR={ 0.5%, 1.0%, 2.0%, 5.0% } [34]. However, since there are few negative aptamers in both datasets, the FPR is 7.1% (Data1) and 8.3% (Data2) when False = 1. So, we used the TPR when FPR is 0 as an alternative metric.

#### 3.4.3 Analysis with ROC curves

The analysis using ROC curve is another way for evaluating binding affinity prediction performance. For the data analysis in section 3.1, the ROC curves for data from rounds 0 to 4 in Data1 are shown in figure S6 and the ROC curves for Data2 are shown in figure S7, respectively. ROC curves corresponding to analysis in section 3.2 are shown in figure S8. In addition, the ROC curves for data for rounds 0 to 8 in Data1 are shown in Figure S9.

Here, ROC curves for data from rounds 0 to 3 in Data1 and round 3 in Data2 are not shown because they were not informative. In these rounds, AME and Enrichment scores were undefined because these scores calculate relative increase of frequency compared to the previous round (Equation 5 and Equation S2) and in these instances either no previous round was present or the sequence was not observed in the previous rounds (Table S6, Table S7). In other words, unless a sequence is observed in successive rounds, AME and Enrichment scores are undefined and the ROC curve is almost a diagonal line. For the same reason, AME and Enrichment values for round 4 data in both Data1 and Data 2 were undefined for some sequences (Figure S6a, Figure S7a).

Besides, since the ROC curves obtained for analysis with RaptRanker and all the existing methods were below the diagonal line for data from rounds 5 to 8 in Data1, we only used data from rounds 0 to 4 for analysis. We also confirmed significant amplification of the unique sequences whose binding-affinity is False data from rounds 5 to 8 in Data1. Frequency, Enrichment, MPBind, and RaptRanker take frequency information into account, so if the False sequences were amplified significantly in the experiment, they are ranked higher. The amplification of the False sequences in Data1, rounds 5–8 might be caused by some experimental errors.

#### 3.4.4 Compare sequence score definitions

As AME is defined as the average motif enrichment of the subsequences from each unique sequence, we can consider best motif enrichment (BME) as the maximum value of motif enrichment. BME of a unique sequence *s* at round *x* is

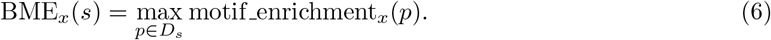

AME assesses the conservation of features over a wider range, whereas BME assesses the conservation of features locally. Since AME is the average, “unique sequences whose nucleotide sequence and secondary structure are conserved as a whole” receive higher scores. In contrast, since BME takes the maximum value, “unique sequences whose nucleotide sequence and secondary are conserved in a specific part” receive higher scores.

We compared AME and BME for the data from the final rounds in both datasets. AME showed higher TPR than BME in both datasets (Table 6). The ROC curve is shown in figure S10. These results indicate that the dataset to be anlysed highly influences the score obtained by either AME or BME. However, AME is more convenient than BME in identifying high-binding affinity sequences because multiple unique sequences that share the same high-score motif have the same BME scores.

**Table 6:**
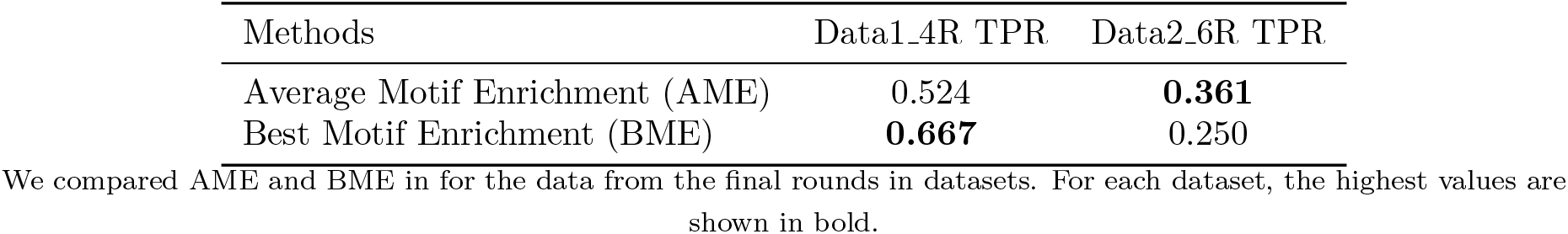
Result of sequence score definitions comparisons

#### 3.4.5 Effect of parameters

We examined the effects changing the values of *wide, weight* and *cosdist* on the performance. We used the parameters (*wide*=10, *weight*=0.5, *cosdist*=0.001, *missing_ratio*=0.00001) on RaptRanker for both data. and observed that *wide*, *weight*, and *cosdist* greatly affected the clustering result. Since the clustering of RaptRanker uses SketchSort, which is an approximation algorithm, we measured the TPR when FPR is 0 for each combination of parameters three times and calculated the average.

The parameters used in this study showed high TPR on average for both datasets, and no significant change in the results was observed when parameters were modified in a range (Supplementary Tables S8 and S9).

#### 3.4.6 Future works

RaptRanker calculates scores for each round and each unique sequence. As a result, RaptRanker, just like the existing methods, cannot detect *bad* rounds (such as rounds 5–8 in Data1). Since RaptRanker uses data from all rounds as the input, we are planning to realize identification of high binding-affinity aptamers in the presence of *bad* rounds.

In the clustering method of RaptRanker, features of subsequences are transformed into sub-sequence-structure profiles (SSSPs), which are clustered using SketchSort and MSF. Since SSSPs need only a matrix of real values, various extensions are possible. For example, RaptRanker can identify DNA aptamers by using DNA secondary structure prediction software instead of CapR. Alternatively, RaptRanker can add new features like G-quadruplex [35] by adding new rows to SSSPs. Generally, this clustering technique can be applied to identification of nucleic acid motifs not only in aptamers but also in non-coding RNAs and/or messenger RNAs. In particular, compared to general k-mer analysis, it is more effective in searching long motifs.

In this study, conventional secondary structures were utilized; however, by considering tertiary structures, which play more important roles in target recognition, in the RaptRanker algorithm, its accuracy could be improved. There are many studies about tertiary structure predictions of RNAs [36, 37, 38], and our future work includes the development of an efficient method to incorporate tertiary structure information into RaptRanker.

## 4 CONCLUSION

We developed RaptRanker, a novel *in silico* method for identifying high binding-affinity aptamers from HT-SELEX data based on local sequence and structural information. RaptRanker determines unique sequences by analyzing data obtained in *all* HT-SELEX rounds, and clusters all subsequences of unique sequences based on similarity in both nucleotide sequence and secondary structure features. Then, RaptRanker identifies high binding-affinity aptamers by calculating AME, which is a score for each unique sequence based on inclusion of clusters.

To evaluate the performance of RaptRanker in identifying high binding-affinity aptamers, we performed two new HT-SELEX experiments, and evaluated sequence sets that include aptamers with low binding-affinity as detected by surface plasmon resonance (SPR) assay. The sequence data which includes the intermediate rounds and evaluation sequence sets are freely available, which will be useful in future aptamer researches.

In both HT-SELEX datasets, RaptRanker showed the best performance in identifying high binding-affinity sequences among all methods tested: Frequency, Enrichment, and MPBind. Moreover, we confirmed the effectiveness of including both nucleotide sequence and secondary structure data in HT-SELEX analysis by examining the identification accuracy of different approaches based on similarity in (i) nucleotide sequence only, (ii) secondary structure only, or (iii) both parameters. In addition, we confirmed that RaptRanker correctly predicts the essential subsequence in each identified sequence. RaptRanker is particularly useful when the existing methods analyzing sequence similarity only cannot provide sufficient identification.

## 5 Data Availability

An implementation of the proposed method, RaptRanker, is available on GitHub https://github.com/hmdlab/RaptRanker.

The two newly acquired HT-SELEX data is available on DRA009383 and DRA009384.

## 6 Funding

This work was supported by JST CREST Grant Number JPMJCR1881, Japan.

## 7 ACKNOWLEDGEMENTS

AY is currently working at a private company in Japan. Computation for this study was partially performed on the NIG supercomputer at ROIS National Institute of Genetics. RI, AY and MH thank Tsukasa Fukunaga and members of Hamada Laboratory for valuable comments. We are also grateful to Satoko Yamazaki, Hisako Ikeda and Hisayo Yasumoto for their early contributions in obtaining part of experimental data. We also thank three anonymous reviewers whose comments greatly improved this manuscript.

## 8 Conflict of interest statement

None declared.

## Notes

### Competing Interest Statement

The authors have declared no competing interest.

### Summary of Updates

1) We included the SPR assay results and transglutaminase assay results in Supplementary Tables S4&S5 and Supplementary Figures S1--S5, which were newly added in this revision. 2) We changed the evaluated sequence set. The new version takes into account differences between batches of experiments, and it is more reliable than the previous one. . Due to this, some parts in our result were changed, which did not affect our previous conclusion. (Please see Tables 2,3,6 Supplementary Figures S6--S10, and Supplementary Tables S8,S9) 3) The new HT-SELEX datasets are now available on DRA009383 and DRA009384. (In the previous submission the upload was completely done, but publishing was not.) We believe these data will be useful in aptamer research.

