## Supplementary Information for "RaptRanker: *in silico* RNA aptamer selection from HT-SELEX experiment based on local sequence and structure information"

#### A Notations

Table S1 summarizes notations frequently utilized in the main manuscript.

Table S1: Notations utilized in the main text

| notation | description |
|---|---|
| $S$ | Set of all unique sequences (in all rounds) |
| $s$ | Each unique sequence, $s \in S$ |
| $P$ | Set of all subsequences |
| $p$ | Each subsequence, $p \in P$ |
| $D_s$ | Set of subsequences from the unique sequence $s$ |
| $\mathfrak{C}$ | Set of all clusters |
| $C$ | Each cluster which is a set of subsequence, $C \in \mathfrak{C}$ |

#### B Calculation of Minimum Spanning Forest

##### B.1 Minimum Spanning Forest (MSF)

Minimum Spanning Forest (MSF) is a weighted unconnected graph in which each connected component becomes a Minimum Spanning Tree (MST). The weighted unconnected graph is one type of graph in which the edges have weights, and not all vertices are necessarily connected. In contrast, a graph in which all vertices must be connected with weights is called weighted connected graph. MST is one type of weighted connected graph in which the sum of the weights of the edges is minimum, and which does not have any closed path.

##### B.2 Kruskal's algorithm

Kruskal's algorithm is an algorithm for calculating MSF. Kruskal's algorithm calculates MSF by using only the edges which do not form a closed path in ascending order of weight. That is, use only the edges whose two vertices belong to different trees (Algorithm S1).

Generally, the time complexity of Kruskal's algorithm is $O(|E| \log |E|)$ for the set of edges $E$. However, the time complexity can be reduced to $O(|E|a(|V|))$ by sorting $E$ in ascending order of its weight beforehand, and using a data structure called UnionFind. $a(n)$ is the inverse function of Ackermann function, which is a very slow increasing function.

##### B.3 UnionFind

UnionFind is one type of disjoint-set data structure. Disjoint-set data structure holds a set of data divided into tree structures in which elements do not overlap each other. UnionFind can perform the following two operations rapidly.

*To whom correspondence should be addressed. Department of Electrical Engineering and Bioscience Faculty of Science and Engineering, Waseda University 55N-06-10, 3-4-1, Okubo Shinjuku-ku, Tokyo 169-8555, Japan. Tel: +81 3 5286 3130; Fax: +81 3 5286 3130;

**Algorithm S1** Kruskal's algorithm

**Require:** A set of edges $E = \{e\}$
**Ensure:** Minimum Spanning Forest $F$ of graph $G(E, V)$ ($V = \{v\}$ is a set of vertices)

```
F = ∅
while E ≠ ∅ do
  Let edge e = {u, v} whose weight is minimum in E
  Remove e from E
  if Vertices u and v belong to different trees in F then
    Add e to F
  end if
end while
```

- Union: Unite two trees into one tree.
- Find: Find the tree to which the element belongs.

The amortized execution time of each operation can be reduced to $a(n)$ by the following contrivances. $a(n)$ is the inverse function of Ackermann function, which is a very slow increasing function.

- Union: Unite the trees where the larger tree as root.
- Find: Replace all references to the parent node which is traced recursively to the references to the root node.

### C Filtering Parameters Used in the Analysis

Table S2: Filtering parameters for Data1

| | |
|---|---|
| *forward_primer* | TAATACGACTCACTATAGGGAGCAGGAGAGAGGTCAGATG |
| *reverse_primer* | CCTATGCGTGCTAGTGTGA |
| *minimum_length* | 25nt |
| *maximum_length* | 35nt |
| *add_forward_primer* | GGGAGCAGGAGAGAGGTCAGATG |
| *add_reverse_primer* | CCTATGCGTGCTAGTGTGA |

The parameters used in the analysis of Data1. The function of each parameter is as follows. *forward_primer*, *reverse_primer* : RaptRanker extracts only the sequences whose both primer binding regions are exactly match these parameters. *minimum_length*, *maximum_length* : RaptRanker extracts only the random regions whose length $l$ is *minimum_length* $\leq l \leq$ *maximum_length*. *add_forward_primer*, *add_reverse_primer* : The secondary structure is predicted to each unique sequence which include these sequences to consider the effect of the primer binding region on the secondary structure.

Table S3: Filtering parameters for Data2

| | |
|---|---|
| *forward_primer* | TAATACGACTCACTATAGGGAGAACTTCGACCAGAAG |
| *reverse_primer* | TATGTGCGCATACATGGATCCTC |
| *minimum_length* | 35nt |
| *maximum_length* | 45nt |
| *add_forward_primer* | GGGAGAACTTCGACCAGAAG |
| *add_reverse_primer* | TATGTGCGCATACATGGATCCTC |

The parameters used in the analysis of Data2. The function of each parameter is as follows. *forward_primer*, *reverse_primer* : RaptRanker extracts only the sequences whose both primer binding regions are exactly match these parameters. *minimum_length*, *maximum_length* : RaptRanker extracts only the random regions whose length $l$ is *minimum_length* $\leq l \leq$ *maximum_length*. *add_forward_primer*, *add_reverse_primer* : The secondary structure is predicted to each unique sequence which include these sequences to consider the effect of the primer binding region on the secondary structure.

### D Calculation of Frequency and Enrichment

#### D.1 Frequency

Frequency value is calculated for each unique sequence $s(\in S)$ and for each round by dividing the number of occurrences of a unique sequence $s$ in a particular round by the number of occurrences of all unique sequences in that round. Frequency of a unique sequence $s$ at round $x$ is

$$\text{Frequency}_x(s) = \frac{\text{count}_x(s)}{\sum_{s' \in S} \text{count}_x(s')} \tag{S1}$$

where $\text{count}_x(s)$ is the occurrence of the sequence $s$ at round $x$.

##### D.2 Enrichment

RaptRanker calculates Enrichment value for each unique sequence $s(\in S)$ and for each round by dividing the Frequency value in a round by the Frequency value in the previous round. Enrichment of a unique sequence $s$ at round $x$ is

$$\text{Enrichment}_x(s) = \frac{\text{Frequency}_x(s)}{\text{Frequency}_{x-1}(s)} \tag{S2}$$

for $\text{Frequency}_{x-1}(s) \neq 0$ and round $x$ is not the first round of inputs; The score is undefined for the other cases.

#### E Evaluated sequence sets

Table S4: Evaluated sequence sets for Data1

| ID | Sequence | Acquisition date | RU | Positive/Negative (1/0) |
|---|---|---|---|---|
| Data1-1 | GTCGAGATTCTGAGGGTTCTCCTGGTGGCT | 161007 | 340.2 | 1 |
| Data1-2 | TGTAGGGAATGACCGGGGTCCTTAACGCTT | 161007 | 315.7 | 1 |
| Data1-3 | TGTAGGGAATGACCGGGTTCCTGGTGCCCG | 161007 | 277.0 | 1 |
| Data1-4 | AACGAGAGATGGTAGGCCTCAAATACAGTC | 161007 | 244.6 | 1 |
| Data1-5 | AACGAGAGATGGTAGGCCTCGCCTTCTAAA | 161007 | 211.3 | 1 |
| Data1-6 | TTAGGAGTTACGAGAGATGTGGCATGTGA | 161007 | 210.5 | 1 |
| Data1-7 | GTACGAGATTCTGAGGGTTCTTTTTGCAC | 161007 | 134.7 | 1 |
| Data1-8 | GTACTTTAGACAACTGAGGGTTCTCTTATT | 161007 | 134.1 | 1 |
| Data1-9 | AACAAGAGATGGTAGGCCTACGTAATGTTT | 161007 | 127.8 | 1 |
| Data1-10 | AACGAGAGATGGTAGGCCTTTGATGATGCT | 161007 | 127.4 | 1 |
| Data1-11 | GCGGAGATTCTGAGGGTTCTCCTGTTCCAG | 161007 | 115.9 | 1 |
| Data1-12 | ACACACGAGAGATGTTGCCTGCTACCCACA | 161007 | 105.5 | 1 |
| Data1-13 | GTCGAGATTTTGAGGGTTCTCCGCTCCAGG | 161007 | 85.1 | 1 |
| Data1-14 | AACAAGAGATGGTAGGCCTAACCATGCTTT | 161007 | 82.2 | 1 |
| Data1-15 | GTCGAGACTTTGAGGGTTCTCCTGCTGTAA | 161007 | 59.8 | 1 |
| Data1-16 | TTAGGAGTTACGAGAGATGTGGCATGTG | 161007 | 51.8 | 1 |
| Data1-17 | ATTAGAAAATTACAAGAGATGTGATCGCTGA | 161007 | 46.8 | 1 |
| Data1-18 | ATACGAGAGATGTAACCCGTCCTGCAATCA | 161007 | 46.2 | 1 |
| Data1-19 | AGGGGGCAAATTGGAAACCCGGGGTTCTAA | 161007 | 44.6 | 1 |
| Data1-20 | AACGAGAGATGGTAGGCCTCCAAGCTCACG | 161007 | 40.4 | 1 |
| Data1-21 | TTTAGGTATTACGAGAGATGTGGATATCGA | 161007 | 38.1 | 1 |
| Data1-22 | ATACGAGAGATGTAGCCTCTTTTATTGCGT | 161007 | 34.4 | 1 |
| Data1-23 | ACACACGAGAGATGTTGCCTGCTACCCGCA | 161007 | 0.7 | 0 |
| Data1-24 | TTTAGTATATACGAGAGATGTGAACAATGA | 161007 | -4.5 | 0 |
| Data1-25 | AAACGAGAGATGTAGCCTTTCTCGAATTCA | 161007 | -7.7 | 0 |
| Data1-26 | TTAGATAATTACAAGAGATGTGGCATCGGA | 161007 | -8.2 | 0 |
| Data1-27 | AACGGTAAACGTGCAAGCCAGTCCTGTTCT | 161007 | -10.3 | 0 |
| Data1-28 | AACAATAAAAATTGTGCAAGCCTTAGGCTC | 161007 | -13.4 | 0 |
| Data1-29 | AACGGTAAACGTGCAAGCCTTGGTAACGTT | 161007 | -14.1 | 0 |
| Data1-30 | AACAATGTAAAATTGTGCAAGCCTATCTT | 161007 | -24.1 | 0 |

The list of evaluated sequence sets of Data1. Sequences are presented in descending order of the RU value. The RU values at the end of protein injection were used for the binding level of aptamers (Figure S1a).

Table S5: Evaluated sequence sets for Data2

| ID | Sequence | Acquisition date | RU | Positive/Negative (1/0) |
|---|---|---|---|---|
| Data2-1 | GCTGTGTCTACGTCCGGATTGGGGACCTGCACGGCCCATG | 170314 | 260.1 | 1 |
| Data2-2 | ACACATAACACGCCTTTGGAGCGATACCTCCATCAACGAT | 170314 | 239.0 | 1 |
| Data2-3 | TCTCCTAGTCCATTTTGGCGCACGCGAATACGGGAGACAC | 170314 | 155.8 | 1 |
| Data2-4 | TAGCTCCGCGCGCACGTTACGCGAAAACGTTCTGCCCGCC | 170314 | 153.0 | 1 |
| Data2-5 | TCCTTAACTTCAGCGTACATTACGCTACATCTCCCCAGCC | 170314 | 120.2 | 1 |
| Data2-6 | TTTCCTAGTCCATTTTGGCGCACGCGAATACGGGAGACAC | 170314 | 113.6 | 1 |
| Data2-7 | AAGGATACGCATCTAGCACCCCTGCCTTCCTTCTAATGTT | 170314 | 109.2 | 1 |
| Data2-8 | TCAATCCTGCACGTACTACGCGAAAACTGTCCCCCGCCTG | 170314 | 102.9 | 1 |
| Data2-9 | TGGAACTACGCGACAAACTTCCCGCCTGTTCCACCTTATG | 170314 | 100.3 | 1 |
| Data2-10 | GTACTACGCCACAACGAACCCCGGCCTGTACCTTTCCGTA | 170314 | 99.7 | 1 |
| Data2-11 | GACTACGCGAAACTTACTCCCCCCGCCTGTCCTTAAACTT | 170314 | 95.7 | 1 |
| Data2-12 | AAACCAGGAGGACGCACATTACGCAAAAAGTATCCCTGCC | 170314 | 92.2 | 1 |
| Data2-13 | TTGCATTCCTGCGCCACATTACGCGAAACGATCCCCGCC | 170323 | 90.8 | 1 |
| Data2-14 | TAGATGTCTTAACGCACATTACGCAAACACACCTCCTGCC | 170323 | 88.5 | 1 |
| Data2-15 | CACTACGTGACCATGATTCCCCACCTGTGCTTCAGTCAGT | 170314 | 85.4 | 1 |
| Data2-16 | GACATCTACGCAAAAGTGTTCCCCTGCCTGATGTGACAAA | 170314 | 83.2 | 1 |
| Data2-17 | TCGGAAGCGATCCGAACATTACGCAAAACACCTCCCTGCC | 170323 | 75.1 | 1 |
| Data2-18 | AGCAAGTACGCGAAAATCTTCCCCGCCTCTTGTTGGCATC | 170314 | 73.9 | 1 |
| Data2-19 | TCACACCCTCTAACGCACATTACGCTAAACCCTCTCAGCC | 170314 | 73.5 | 1 |
| Data2-20 | AATGCTAAAATCCAGCGGCGCACGCGATGACGGGAGAAAC | 170323 | 73.0 | 1 |
| Data2-21 | TCTAGACACTACGCGAAAAGAACCCACGCCTGTGTCTACG | 170314 | 65.4 | 1 |
| Data2-22 | TGCCAAGCCAGGCAGCGTGCATAATCCGAGTAGACATTTT | 170314 | 65.1 | 1 |
| Data2-23 | ACAAATATGCAACATGTTACGCCAAAAGTTCCCCGGCCTA | 170314 | 62.6 | 1 |
| Data2-24 | TTATCACCCCTGCACATTACGCAAAAGTATTATCCCTGCC | 170314 | 57.2 | 1 |
| Data2-25 | TGCCATAAGCATACGCAACAAACTCGCCTGCCTTGCTTAG | 170323 | 55.5 | 1 |
| Data2-26 | AAAGACGGTACGCGACTCAACACCCGCCTCCGTCAGGTTT | 170314 | 52.3 | 1 |
| Data2-27 | GTGCACGGCAATACGCATGACGTTCACCCTGCCTTTGCCT | 170314 | 51.1 | 1 |
| Data2-28 | TACGACTACTTCGGCACATTACGCGACGATCGAACCCGCC | 170323 | 47.0 | 1 |
| Data2-29 | GTTCAAATCCCCGCACATTACGTAACACTCGAACCCTACC | 170314 | 46.9 | 1 |
| Data2-30 | ATTCCAGAGAGCGACACATTACGCGAAACACTCGCCCGCC | 170314 | 45.8 | 1 |
| Data2-31 | ACTATATCCGCCGTACTACGCATCTTTTTCTTCCTGCCTG | 170314 | 45.2 | 1 |
| Data2-32 | CTGGATTACGCGAAAAAACTCCTCGCCTATCGCAGCAATG | 170314 | 40.6 | 1 |
| Data2-33 | TCCTTGCCACCGTCTCACATTACGCTAAATCTCTCCAGCC | 170323 | 39.1 | 1 |
| Data2-34 | TTCCGGCCTACGCAACTTTTTATCCCCTGCCTGGCCGTAAT | 170314 | 35.8 | 1 |
| Data2-35 | TTCCTGCCAATTGAGCACATTACGCAAAAAAATTCCCTGCC | 170314 | 29.5 | 1 |
| Data2-36 | GAAGACAAGCGCACACAACGGATGACGGGAGAAACAATGA | 170314 | 27.6 | 1 |
| Data2-37 | CACCTCTCCCAACAGCGGAACTGTTTCTCGGTCCAAAAGG | 170314 | 24.8 | 0 |
| Data2-38 | CACTACGCAAACTCCTTCTGCCTGTGCTTGAAAAACGCAT | 170314 | 24.1 | 0 |
| Data2-39 | CTACCAGGTCTTGCACATTACGCGACAACCAAGCCCCGCC | 170314 | 21.9 | 0 |
| Data2-40 | TAGCTCATAACGACCAAAACGACCCAGCAGTTCCTTTCGT | 170314 | 21.6 | 0 |
| Data2-41 | ACCAATAACCGTTCACATTACGCATCAACTCAACCTGCC | 170323 | 18.6 | 0 |
| Data2-42 | TAAGACGCACCATAACTACGCAAGAGTTCTCTCTGCCTGA | 170314 | 18.2 | 0 |
| Data2-43 | TCCACTCTCGCGCACATATACGCAAAACCCTACCTGCCT | 170314 | 15.4 | 0 |
| Data2-44 | TCCTACACTGCGCATAATACGTAACTATCGACCTACCTT | 170314 | 8.8 | 0 |
| Data2-45 | AGCGACTAAACGCAGAGAAATCGTCTCCCTGAGGTTCCGA | 170314 | 5.0 | 0 |
| Data2-46 | TCCTAAGATGCGTTAACATTACGTAAGAATCTACCTACC | 170314 | 3.5 | 0 |
| Data2-47 | ATTGTGACCCGATTTGTCACTCTTTCTATGTACGCGGAAT | 170314 | -0.1 | 0 |
| Data2-48 | GACTACCATCGAACGTACATTACGCTACTCTCTACCAGC | 170323 | -4.2 | 0 |

The list of evaluated sequence sets of Data2. Sequences are presented in descending order of the RU value. The RU values at the end of protein injection were used for the binding level of aptamers (Figure S2a).

### F The results of SPR assay

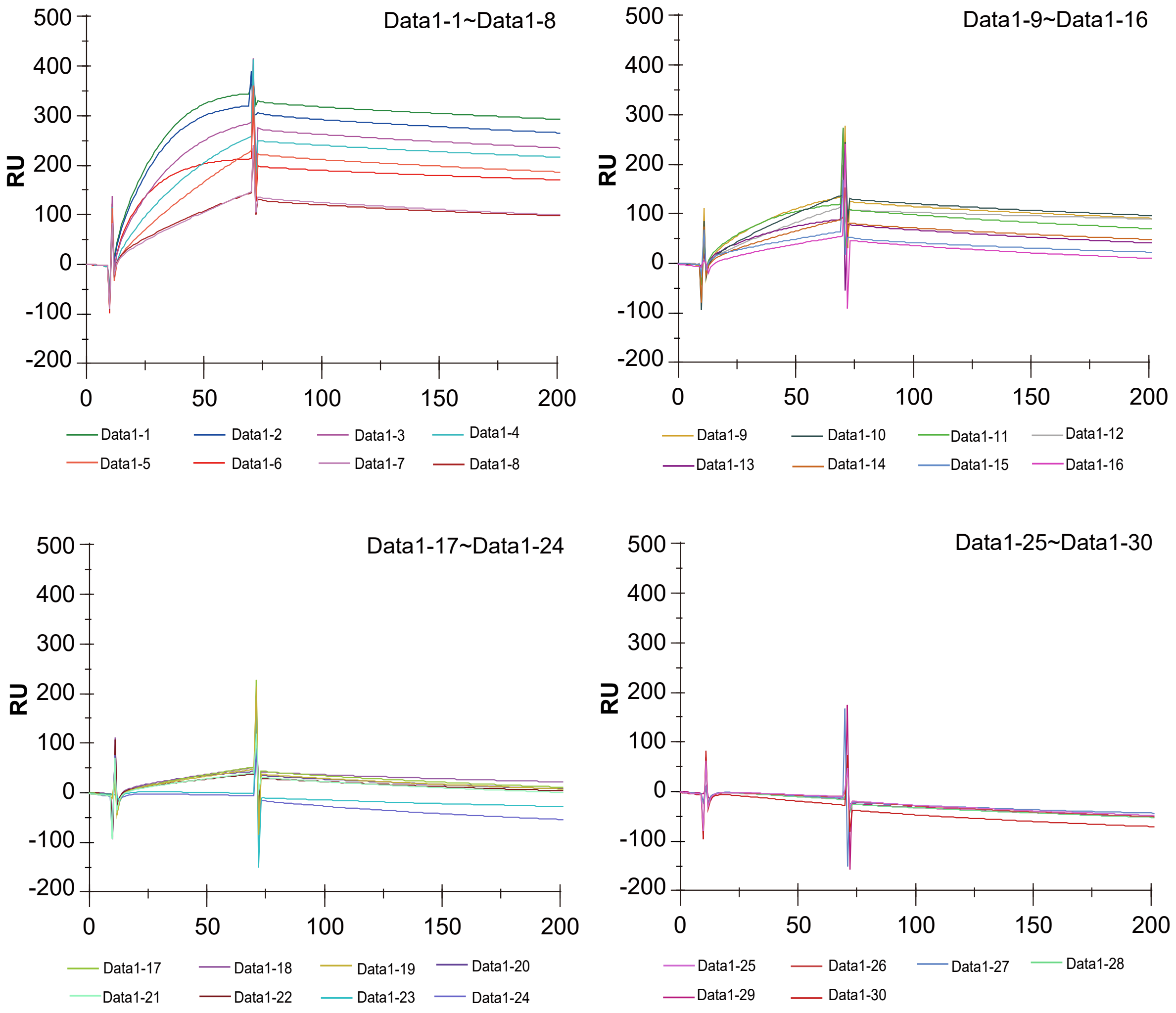


(a)

| Methods | 1st | 2nd | 3rd |
|---|---|---|---|
| Frequency | Data1-2 | Data1-3 | Data1-13 |
| Enrichment | Data1-2 | Data1-4 | Data1-15 |
| MPBind | Data1-24 | Data1-21 | Data1-22 |
| RaptRanker | Data1-3 | Data1-2 | Data1-11 |

(b)

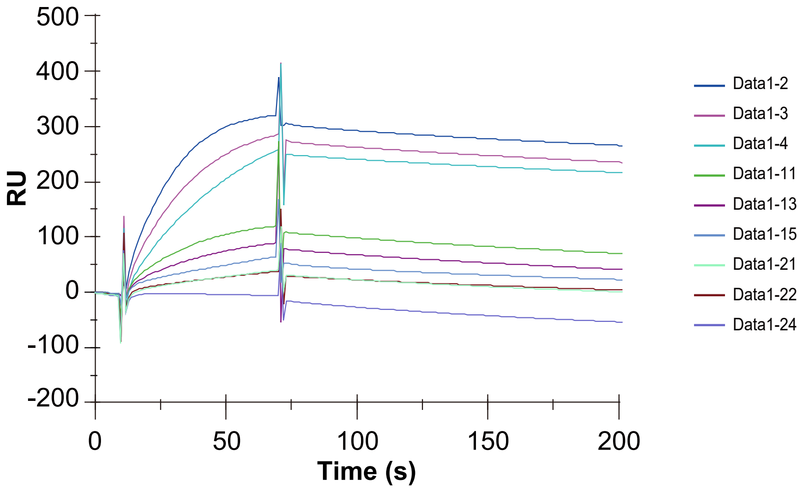


(c)

Figure S1: The figures show SPR sensorgrams of the association and dissociation phases of aptamers. The RU values at the end of protein injection were used for the binding level of aptamers. (a) SPR sensorgrams for all aptamers from the evaluated sequence sets for Data1. (b) List of the top 3 aptamers from each method (Frequency, Enrichment, MPBind, and RaptRanker) used to the evaluated sequence sets for Data1. (c) SPR sensorgrams for the top 3 aptamers from each method used to the evaluated sequence sets for Data1. These sensorgrams are re-displayed from Figure S1a.

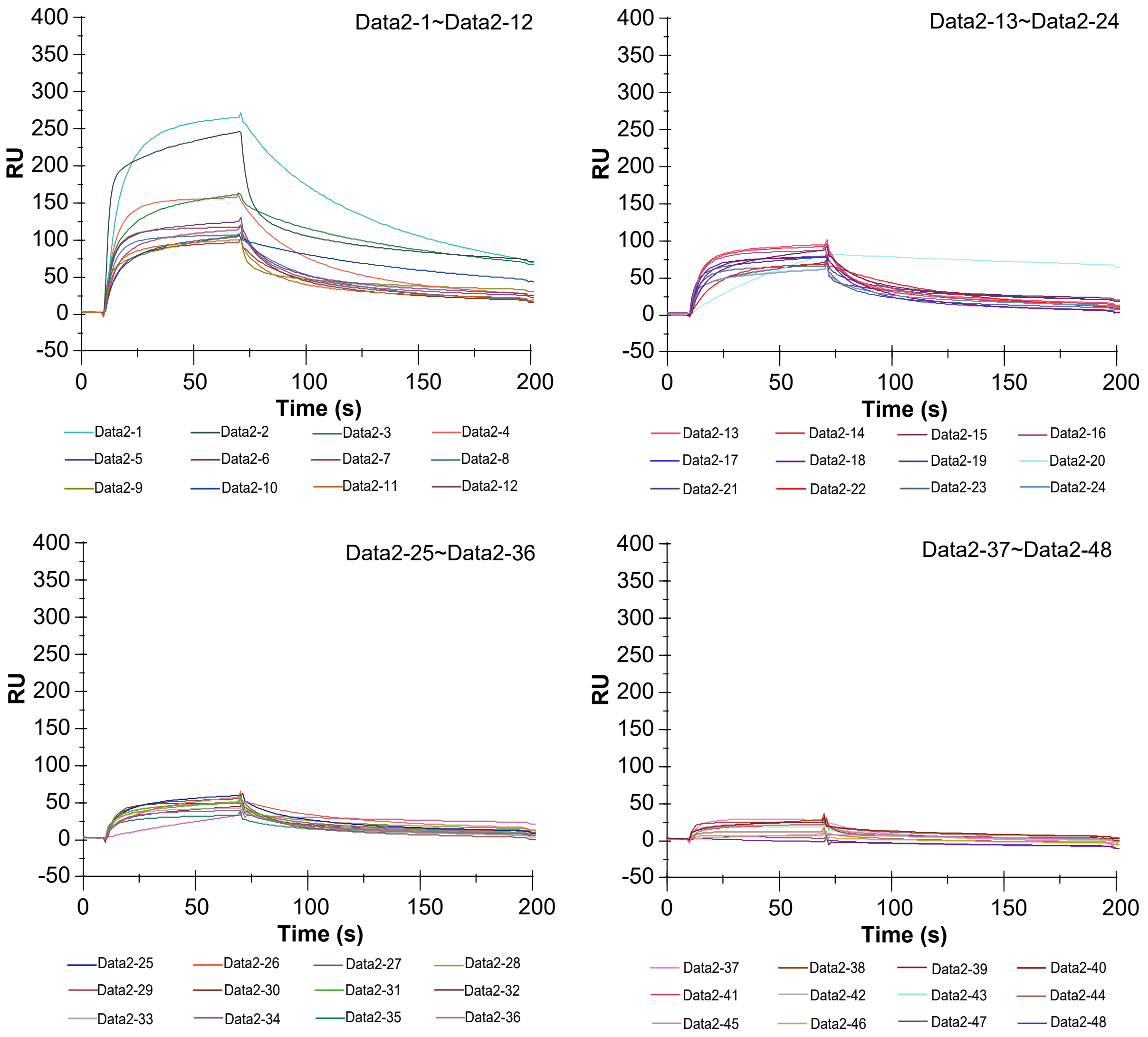


(a)

| Methods | 1st | 2nd | 3rd |
|---|---|---|---|
| Frequency | Data2-10 | Data2-20 | Data2-3 |
| Enrichment | Data2-1 | Data2-20 | Data2-6 |
| MPBind | Data2-3, Data2-10, Data2-16, Data2-20 | | |
| RaptRanker | Data2-1 | Data2-20 | Data2-6 |

(b)

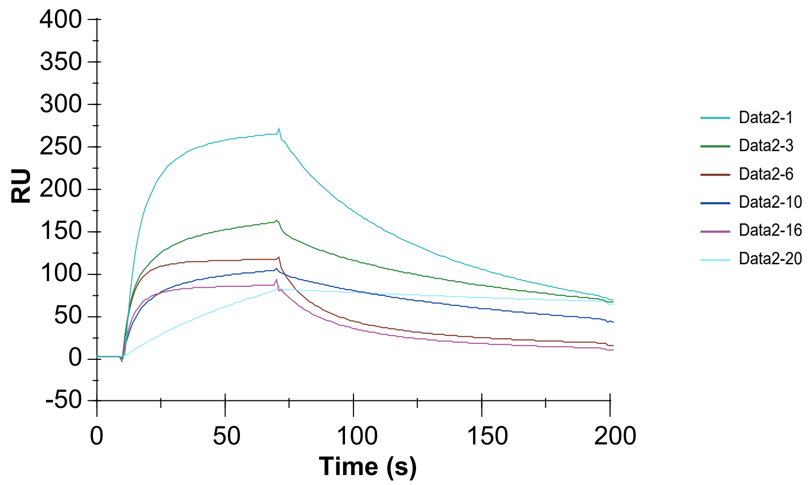


(c)

Figure S2: The figures show SPR sensorgrams of the association and dissociation phases of aptamers. The RU values at the end of protein injection were used for the binding level of aptamers. (a) SPR sensorgrams for all aptamers from the evaluated sequence sets for Data2. (b) List of the top 3 aptamers from each method (Frequency, Enrichment, MPBind, and RaptRanker) used to the evaluated sequence sets for Data2. (c) SPR sensorgrams for the top 3 aptamers from each method used to the evaluated sequence sets for Data2. These sensorgrams are re-displayed from Figure S2a.

### G Truncation of sequences

| sequence | $IC_{50}$ (μM) |
|---|---|
| GGGAGCAGGAGAGAGGUCAGAUGAACGAGAGAUGGUAGGCCUUUGAUGAUGCUCCUAUGCGUGCUAGUGUGA | 0.140±0.016 |
| GGGAGCAGGAGAGAGGUCAGAUGAACGAGAGAUGGUAGGCCUUUGAUGAUGCUCCC | 0.267±0.029 |
| GGGAGGAGAGAGGUCAGAUGAACGAGAGAUGGUAGGCCUUUGAUGAUCCC | 0.243±0.055 |
| GGGAGAGAGGUCAGAUGAACGAGAGAUGGUAGGCCUUUGAUCCC | 0.350±0.098 |
| GGAGAGGUCAGAUGAACGAGAGAUGGUAGGCCUUUCC | 0.167±0.024 |
| GGAGGUCAGAUGAACGAGAGAUGGUAGGCCUCC | 0.185±0.041 |
| GGGGUCAGAUGAACGAGAGAUGGUAGGCCCC | 0.266±0.026 |
| GGGUCAGAUGAACGAGAGAUGGUAGGCCC | 0.231±0.029 |
| GGUCAGAUGAACGAGAGAUGGUAGGCC | 0.189±0.029 |
| GGCAGAUGAACGAGAGAUGGUAGCC | 0.427±0.173 |

GGGAGCAGGAGAGAGGUCAGAUGAACGAGAGAUGGUAGGCCUUUGAUGAUGCUCCUAUGCGUGCUAGUGUGA

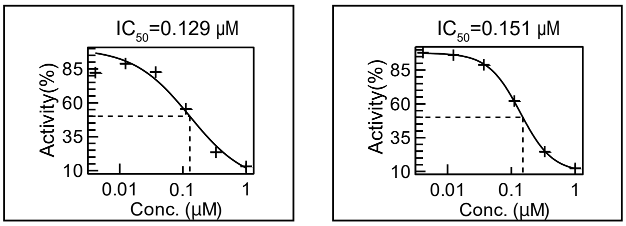


GGGAGCAGGAGAGAGGUCAGAUGAACGAGAGAUGGUAGGCCUUUGAUGAUGCUCCC

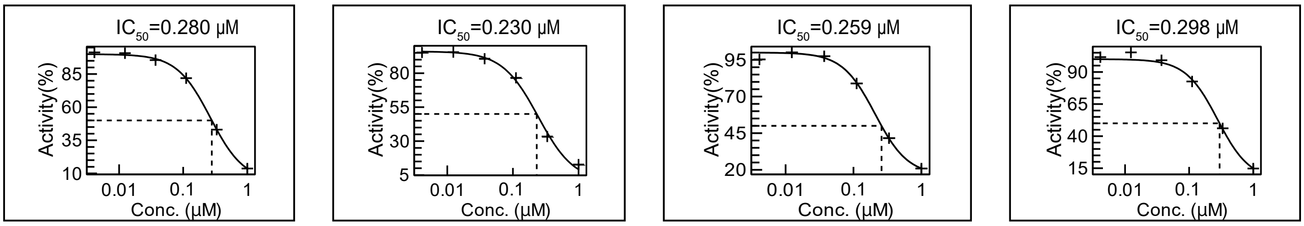


GGGAGGAGAGAGGUCAGAUGAACGAGAGAUGGUAGGCCUUUGAUGAUCCC

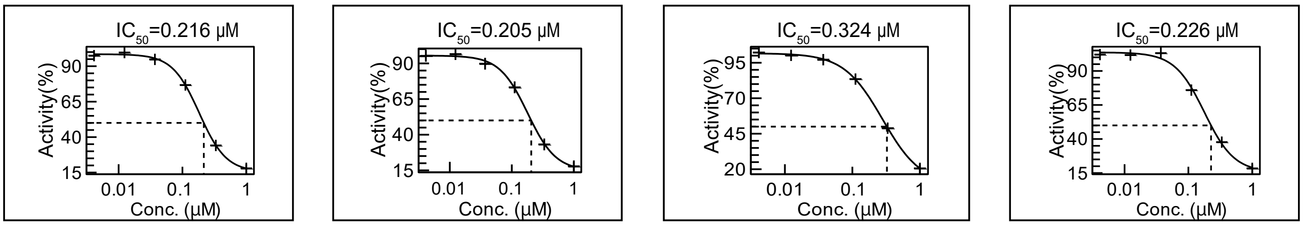


GGGAGAGAGGUCAGAUGAACGAGAGAUGGUAGGCCUUUGAUCCC

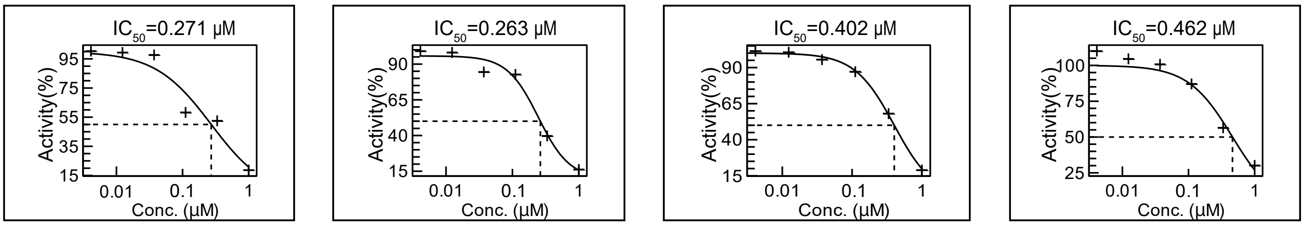

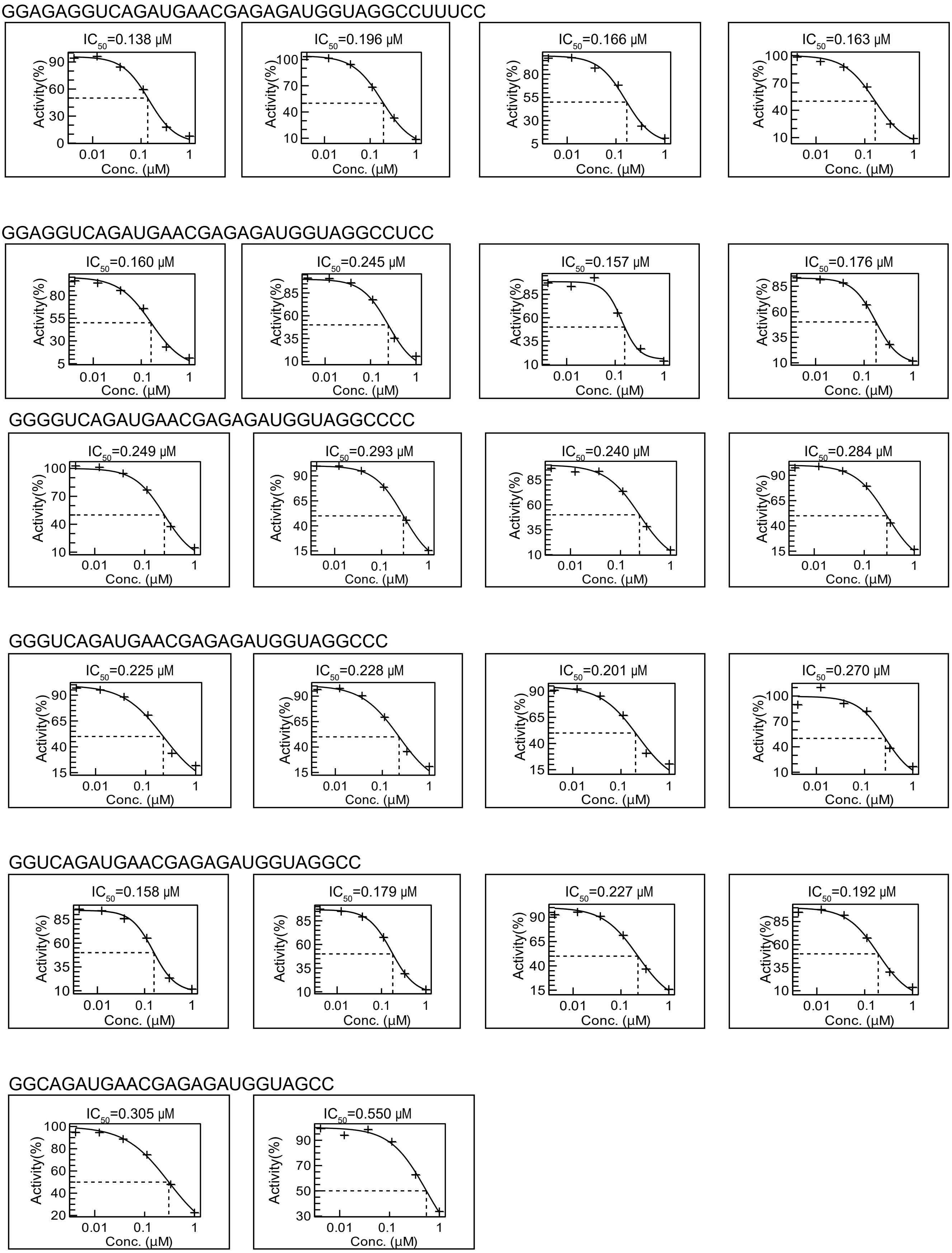


Figure S3: **The truncation of a high binding-affinity aptamer sequence of Data1.** We truncated a high binding-affinity aptamer sequence of Data1 based on the Transglutaminase assay. The sequence in the top row is the original sequence, and the sequence in the bottom row is the minimal active sequence. The figures show TG2 activity in the presence of indicated aptamers. Experimental procedures are described in the "Transglutaminase assay" section. Broken lines indicate half maximal inhibitory concentration ($IC_{50}$).

| ID | sequence | Relative binding (%) |
|---|---|---|
| S2−1 | GGGAGAACUUCGACCAGAAGGCUGUGUCUACGUCCGGAUUGGGGACCUGCACGGCCCAUGUAUGUGCGCAUACAUGGAUCCUC | 100 |
| S2−2 | GGACCAGAAGGCUGUGUCUACGUCCGGAUUGGGGACCUGCACGGCCCAUGUAUGUGCGCAUACAUGGAUCCUC | 19 |
| S2−3 | GGGCUGUGUCUACGUCCGGAUUGGGGACCUGCACGGCCCAUGUAUGUGCGCAUACAUGGAUCCUC | 7 |
| S2−4 | GGGAGAACUUCGACCAGAAGGCUGUGUCUACGUCCGGAUUGGGGACCUGCACGGCCCAUGUAUGUGCGCAUAC | 16 |
| S2−5 | GGGAGAACUUCGACCAGAAGGCUGUGUCUACGUCCGGAUUGGGGACCUGCACGGCCCAUGUAU | 12 |
| S2−6 | GGGAGAACUUCGACCAGAAGGCUGUGUCUACGUCCGGAUUGGGGACCUGCACGGC | 29 |
| S2−7 | GGGCUGUGUCUACGUCCGGAUUGGGGACCUGCACGGC | 9 |

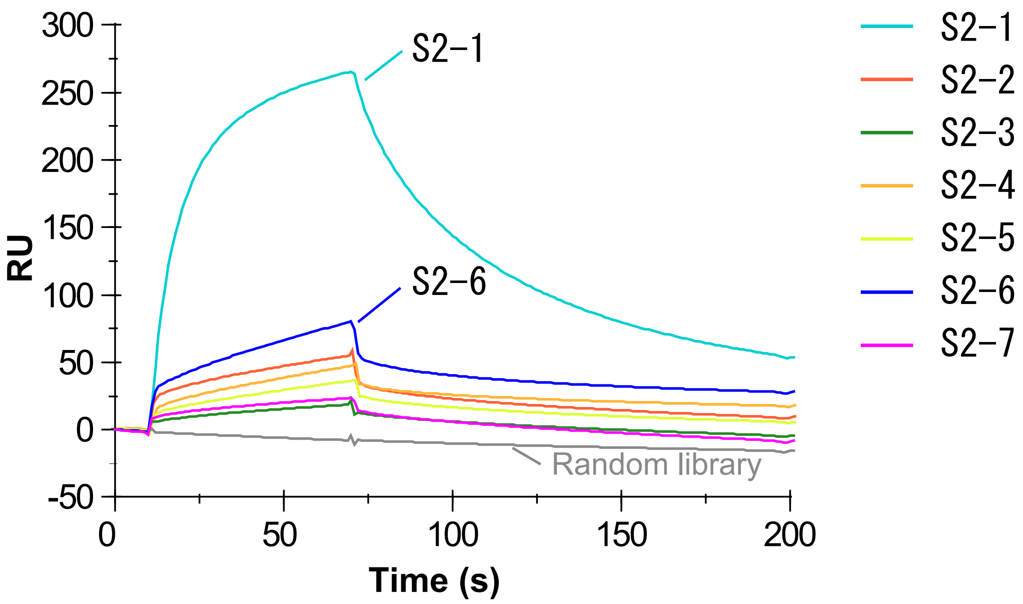


Figure S4: **The truncation of a high binding-affinity aptamer sequence of Data2.** We truncated a high binding-affinity aptamer sequence of Data2 based on SPR assay. In the table, the sequence of S2-1 is the original sequence, and the sequence of S2-7 is the minimal active sequence. The SPR sensorgrams in the bottom figure show the association and dissociation phases for each aptamer.

| ID | sequence | Relative binding (%) |
|---|---|---|
| Mother | GGGAGAACUUCGACCAGAAGGCUGUGUCUACGUCCGGAUUGGGGACCUGCACGGCCCAUGUAUGUGCGCAUACAUGGAUCCUC | 100 |
| S3-1 | GGGAGAACUUCGACCAGAAGGCUGUGU UACGUCCGGAUUGGGGACCUGCACGGCCCAUGUAUGUGCGCAUACAUGGAUCCUC | 139 |
| S3-2 | GGGAGAACUUCGACCAGAAGGCUGUGUC ACGUCCGGAUUGGGGACCUGCACGGCCCAUGUAUGUGCGCAUACAUGGAUCCUC | 1 |
| S3-3 | GGGAGAACUUCGACCAGAAGGCUGUGUCU CGUCCGGAUUGGGGACCUGCACGGCCCAUGUAUGUGCGCAUACAUGGAUCCUC | 0 |
| S3-4 | GGGAGAACUUCGACCAGAAGGCUGUGUCUA GUCCGGAUUGGGGACCUGCACGGCCCAUGUAUGUGCGCAUACAUGGAUCCUC | 0 |
| S3-5 | GGGAGAACUUCGACCAGAAGGCUGUGUCUAC UCCGGAUUGGGGACCUGCACGGCCCAUGUAUGUGCGCAUACAUGGAUCCUC | 0 |
| S3-6 | GGGAGAACUUCGACCAGAAGGCUGUGUCUACG CCGGAUUGGGGACCUGCACGGCCCAUGUAUGUGCGCAUACAUGGAUCCUC | 0 |
| S3-7 | GGGAGAACUUCGACCAGAAGGCUGUGUCUACGU CGGAUUGGGGACCUGCACGGCCCAUGUAUGUGCGCAUACAUGGAUCCUC | 16 |
| S3-8 | GGGAGAACUUCGACCAGAAGGCUGUGUCUACGUCC GAUUGGGGACCUGCACGGCCCAUGUAUGUGCGCAUACAUGGAUCCUC | 182 |
| S3-9 | GGGAGAACUUCGACCAGAAGGCUGUGUCUACGUCCGG UUGGGGACCUGCACGGCCCAUGUAUGUGCGCAUACAUGGAUCCUC | 130 |
| S3-10 | GGGAGAACUUCGACCAGAAGGCUGUGUCUACGUCCGGA UGGGGACCUGCACGGCCCAUGUAUGUGCGCAUACAUGGAUCCUC | 120 |
| S3-11 | GGGAGAACUUCGACCAGAAGGCUGUGUCUACGUCCGGAUU GGGACCUGCACGGCCCAUGUAUGUGCGCAUACAUGGAUCCUC | 174 |
| S3-12 | GGGAGAACUUCGACCAGAAGGCUGUGUCUACGUCCGGAUUGGGG CCUGCACGGCCCAUGUAUGUGCGCAUACAUGGAUCCUC | 43 |
| S3-13 | GGGAGAACUUCGACCAGAAGGCUGUGUCUACGUCCGGAUUGGGGAC UGCACGGCCCAUGUAUGUGCGCAUACAUGGAUCCUC | 0 |
| S3-14 | GGGAGAACUUCGACCAGAAGGCUGUGUCUACGUCCGGAUUGGGGACC GCACGGCCCAUGUAUGUGCGCAUACAUGGAUCCUC | 0 |
| S3-15 | GGGAGAACUUCGACCAGAAGGCUGUGUCUACGUCCGGAUUGGGGACCU CACGGCCCAUGUAUGUGCGCAUACAUGGAUCCUC | 3 |
| | (putative motif) **UACGU** **CUG** | N/A |

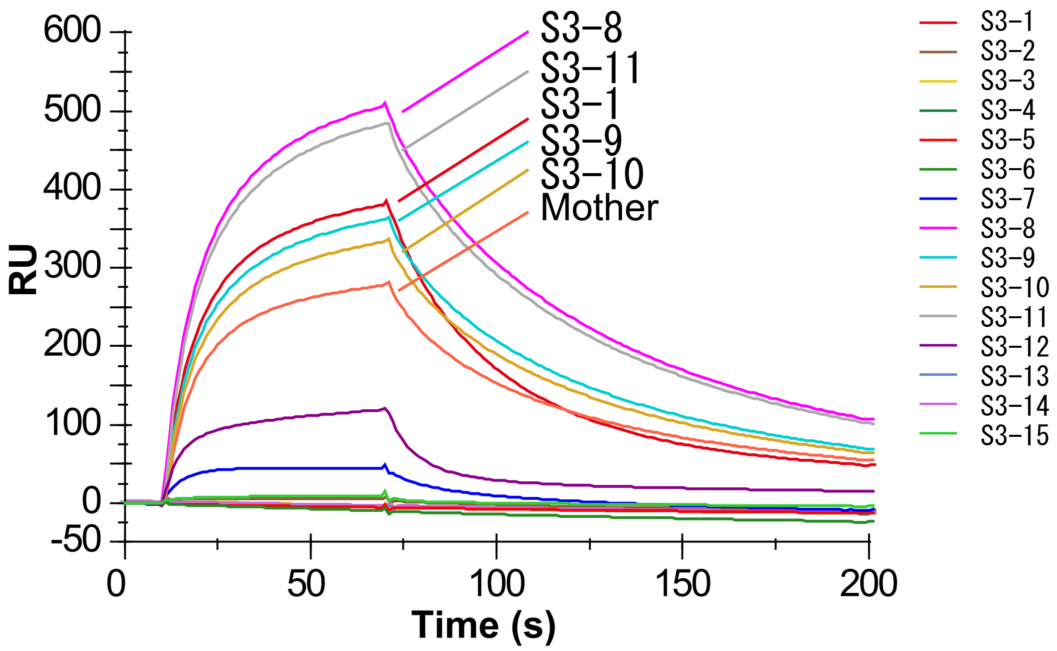


Figure S5: **Binding motif evaluation.** We predict a binding motif of Data2 based on SPR assay for a series of mutant aptamers with single nucleotide deletion (shown in the top Table). The sequence of "Mother" is the original sequence, and the putative motif is shown in the bottom row of the table. The SPR sensorgrams in the bottom figure show the association and dissociation phases for each aptamer.

### H Calculation of Average K-mer Enrichment (AKE)

As a method analyzing only sequence similarity, we calculated AKE in a similar way we calculated AME based on RaptRanker's clustering. Let $\mathfrak{K}$ be the set of all k-mers, and let $K$ be a k-mer. In this case, since $P$ is a set of all subsequences, $\cup_{K \in \mathfrak{K}} K = P$ holds. Also, since k-mers do not overlap each other, $K \cap K' = \emptyset$ holds for any pair of k-mers $K, K' (\in \mathfrak{K})$. AKE is calculated in the same way as AME by replacing the cluster $C$ in equation **2**, equation **3**, equation **4**, and equation **5** with k-mer $K$.

RaptRanker calculates k-mer frequency for each k-mer $K(\in \mathfrak{K})$and for each round by summing up the subsequence frequencies of subsequences that belong to that k-mer. K-mer frequency of a k-mer $K$ at round $x$ is

$$\text{kmer\_frequency}_x(K) = \sum_{p \in K} \text{subsequence\_frequency}_x(p). \tag{S3}$$

Then, RaptRanker calculates k-mer enrichment for each k-mer $K(\in \mathfrak{K})$ and for each round by dividing the k-mer frequency in a round by the k-mer frequency in the previous round. k-mer enrichment of a k-mer $K$ at round $x$ is

$$\text{kmer\_enrichment}_x(K) = \frac{\text{kmer\_frequency}_x(K)}{\text{kmer\_frequency}_{x-1}(K)} \tag{S4}$$

for $\text{kmer\_frequency}_{x-1}(K) \neq 0$ and round $x$ is not the first round; otherwise the score is undefined.

RaptRanker defines k-mer enrichment of each k-mer $K(\in \mathfrak{K})$ as the score of each subsequence belonging to that k-mer $p(\in K)$, such that,

$$\text{kmer\_enrichment}_x(p) = \text{kmer\_enrichment}_x(K) \tag{S5}$$

where $p \in K$.

RaptRanker calculates AKE for each unique sequence $s(\in S)$ and for each round by taking the average of the k-mer enrichment values of subsequences from a unique sequence. The AKE of a unique sequence $s$ at round $x$ is defined as

$$\text{AKE}_x(s) = \frac{\sum_{p \in D_s} \text{kmer\_enrichment}_x(p)}{|D_s|}. \tag{S6}$$

If round $x$ is the first round, the score is undefined.

### I ROC curves

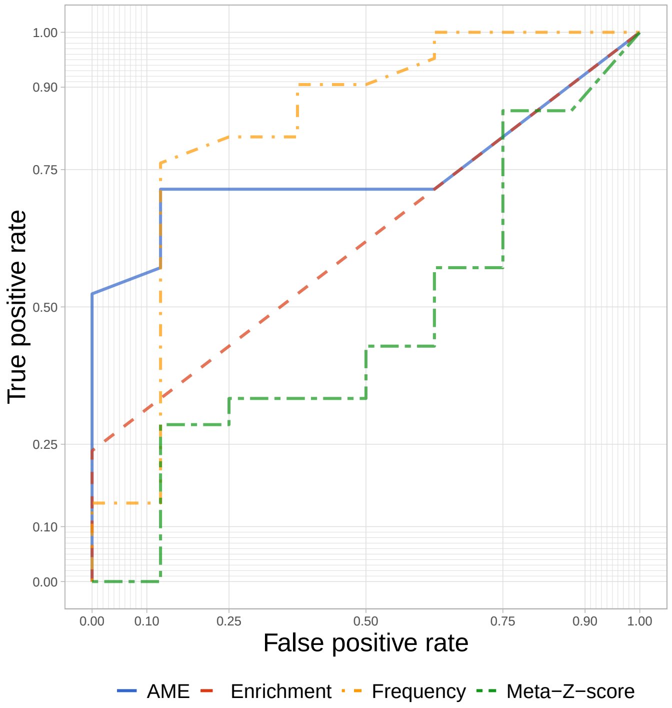


(a) 4R

Figure S6: The ROC curves for Data1 rounds 0–4R. The ROC curves for rounds 0–3R are not listed because they are not informative because for those rounds, AME and Enrichment values of almost all sequences were undefined due to insufficient enrichments of pools .

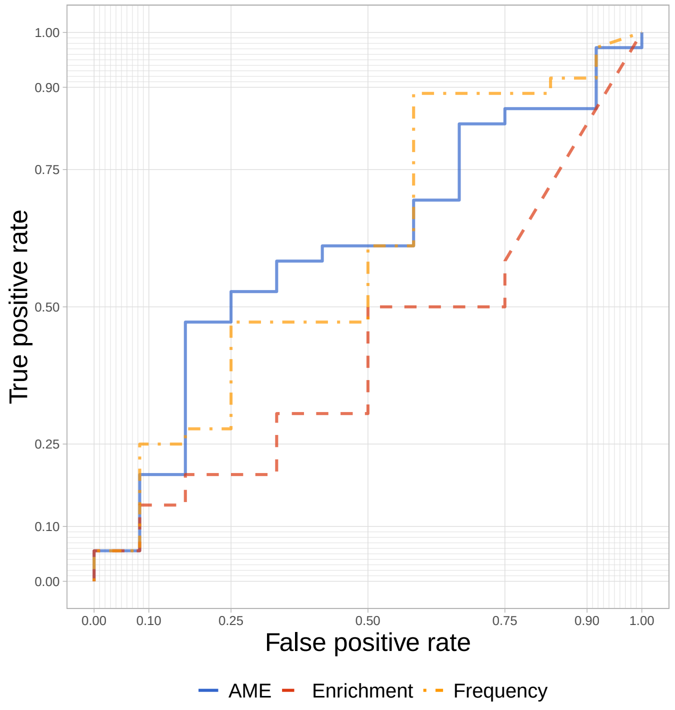


(a) 4R

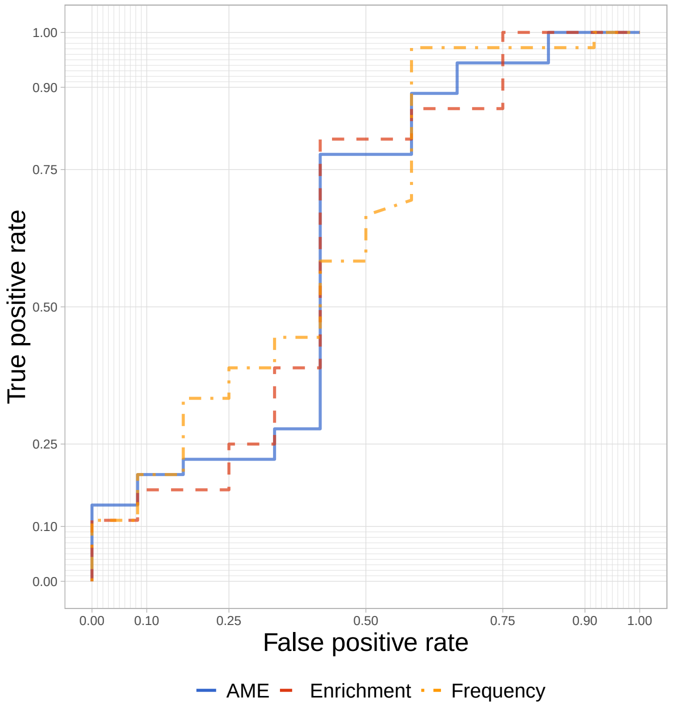


(b) 5R

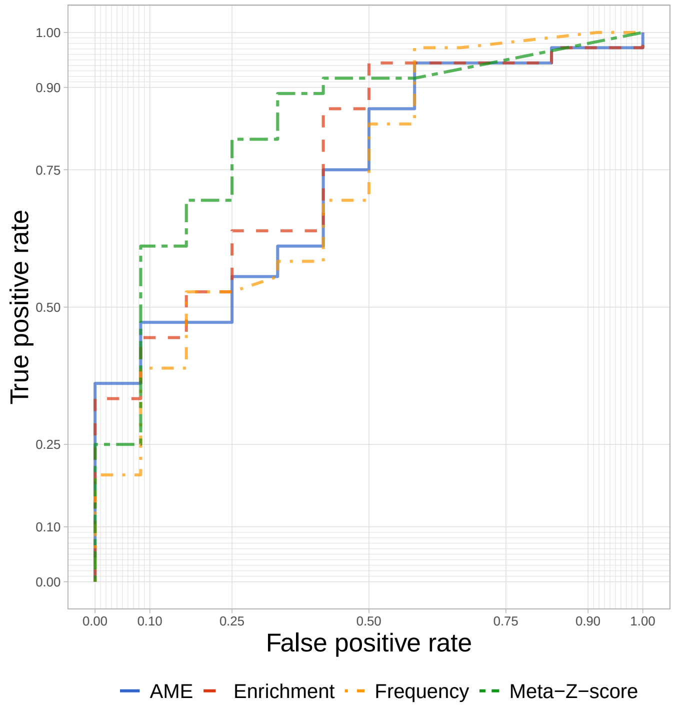


(c) 6R

Figure S7: The ROC curves for Data2. The ROC curves for round 3 are not shown because they were not informative; AME and Enrichment values of all sequences were undefined for round 3 since it was the first round included in the analysis.

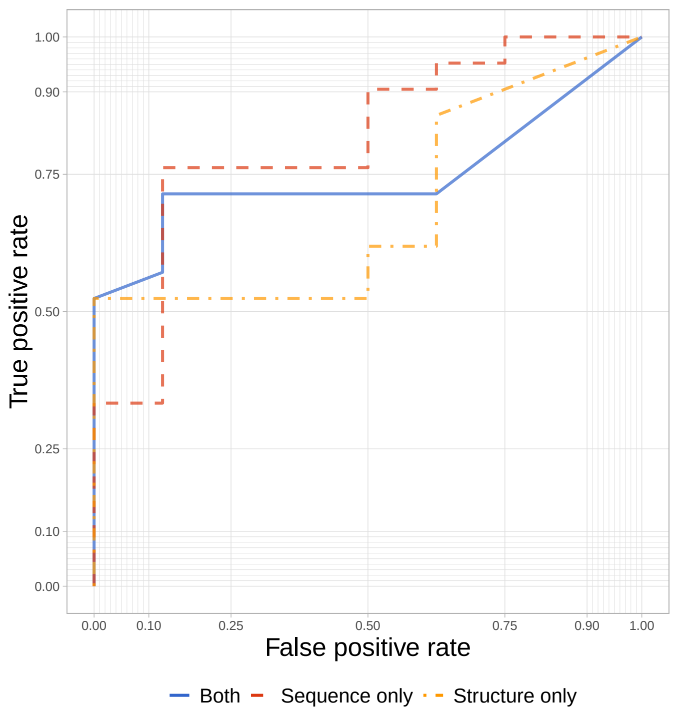


(a) RAPT1_4R

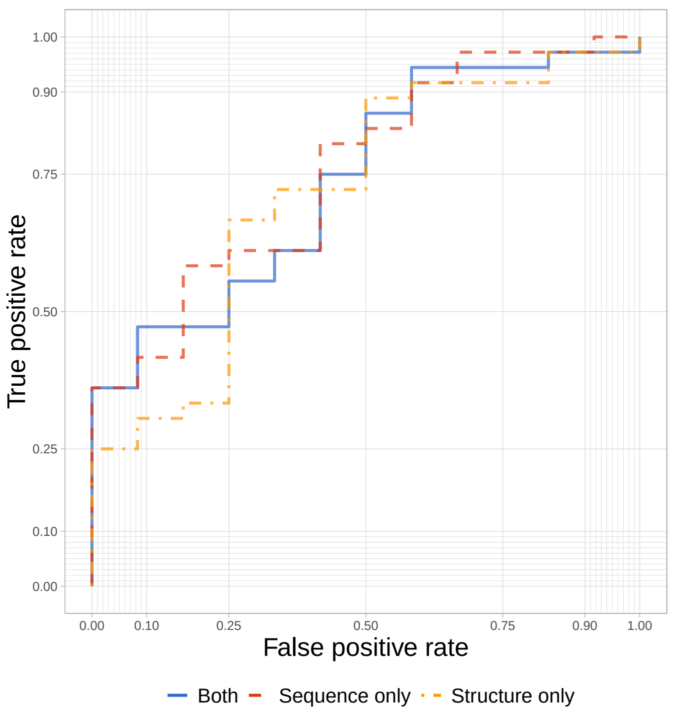


(b) RAPT3_6R

Figure S8: ROC curves corresponding to analysis in section "Validation of the effectiveness of analyzing both sequence and secondary structure". Only the results from rounds 4 and 6 are shown because they are the last rounds of each dataset.

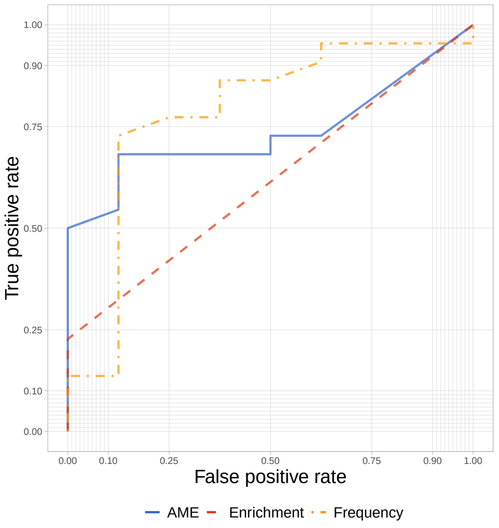


(a) 4R

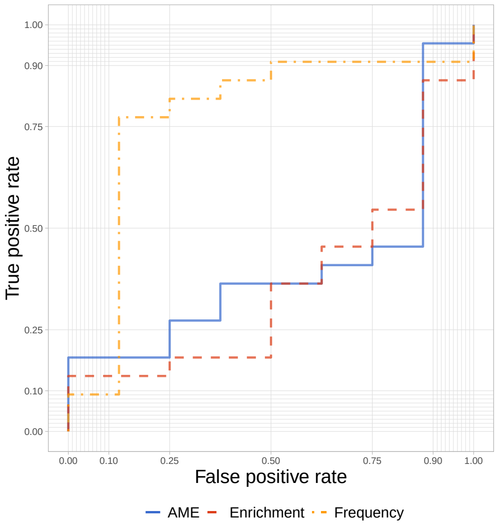


(b) 5R

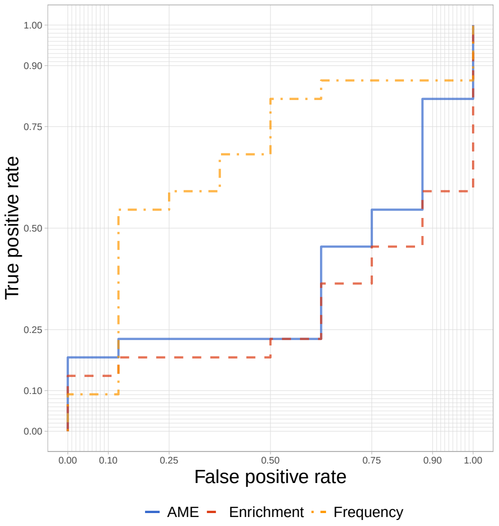


(c) 6R

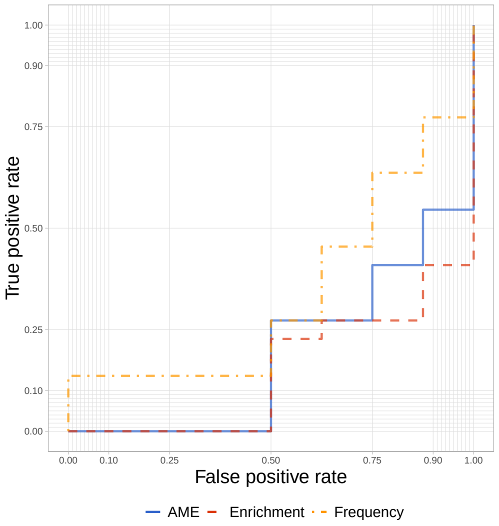


(d) 7R

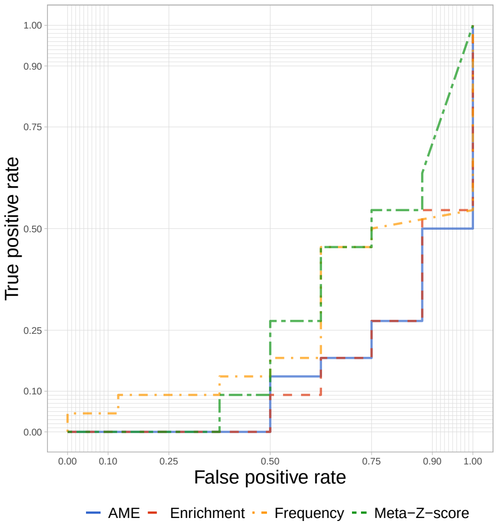


(e) 8R

Figure S9: ROC curves for Data1, rounds 0–8R. The ROC curves for rounds 0–3R were not informative because, for those rounds, AME and Enrichment values of almost all sequences were undefined due to insufficient enrichments of pools.

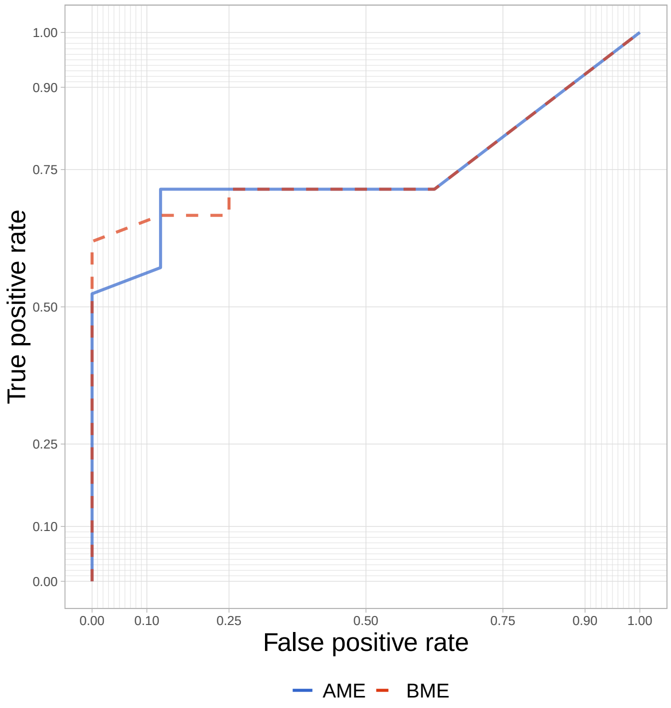


(a) Data1_4R

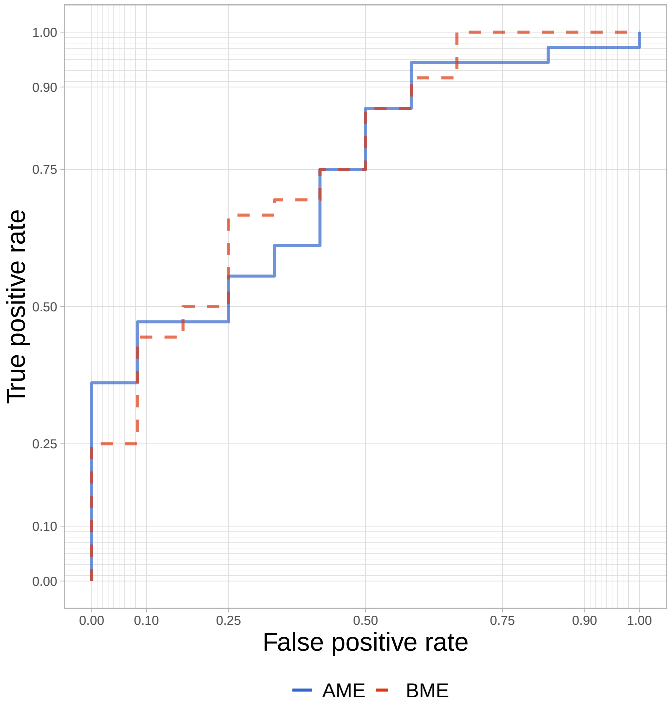


(b) Data2_6R

Figure S10: ROC curves corresponding to analysis in section "Compare sequence score definitions". Only the results from rounds 4 and 6 are shown because they are the last rounds of each dataset.

### J The Number of Sequences in Each Round of Each Dataset

Table S6: The number of sequences in each round of Data1

| round | raw | filtered | unique | filtered/raw | unique/raw | unique/filtered |
|---|---|---|---|---|---|---|
| 0R | 162003 | 114663 | 113106 | 70.78% | 69.82% | 98.64% |
| 1R | 91610 | 72555 | 71453 | 79.20% | 78.00% | 98.48% |
| 2R | 45431 | 36583 | 36071 | 80.52% | 79.40% | 98.60% |
| 3R | 91441 | 72758 | 71719 | 79.57% | 78.43% | 98.57% |
| 4R | 80864 | 64327 | 49260 | 79.55% | 60.92% | 76.58% |
| 5R | 108428 | 86805 | 28582 | 80.06% | 26.36% | 32.93% |
| 6R | 98237 | 80920 | 11825 | 82.37% | 12.04% | 14.61% |
| 7R | 49469 | 40341 | 5092 | 81.55% | 10.29% | 12.62% |
| 8R | 113137 | 81161 | 6029 | 71.74% | 5.33% | 7.43% |
| to 4R total | 471349 | 360886 | 341415 | 76.56% | 72.43% | 94.60% |
| to 8R total | 840620 | 650113 | 379534 | 77.34% | 45.15% | 58.34% |

Titles "raw", "filtered", "unique" indicate the number of sequences inputted, the number of sequences to be analyzed, and the number of unique sequences, respectively.

Table S7: The number of sequences in each round of Data2

| round | raw | filtered | unique | filtered/raw | unique/raw | unique/filtered |
|---|---|---|---|---|---|---|
| 3R | 146505 | 98861 | 96491 | 67.48% | 65.86% | 97.60% |
| 4R | 121185 | 80218 | 5889 | 66.19% | 4.86% | 7.34% |
| 5R | 116917 | 80114 | 6954 | 68.52% | 5.95% | 8.68% |
| 6R | 82488 | 47360 | 4813 | 57.41% | 5.83% | 10.16% |
| total | 467095 | 306553 | 169924 | 65.63% | 36.38% | 55.43% |

Titles "raw", "filtered", "unique" indicate the number of sequences inputted, the number of sequences to be analyzed, and the number of unique sequences, respectively.

### K Effect of parameters

Table S8: Effect of parameters on Data1 (0R to 4R)

| *cosdist* | *wide* | *weight* | repeat_1 | repeat_2 | repeat_3 | Average TPR |
|---|---|---|---|---|---|---|
| 1.00E-04 | 8 | 0.1 | 0.524 | 0.524 | 0.524 | 0.524 |
| | | 0.5 | 0.524 | 0.524 | 0.524 | 0.524 |
| | | 1 | 0.619 | 0.619 | 0.619 | 0.619 |
| | 10 | 0.1 | 0.238 | 0.238 | 0.238 | 0.238 |
| | | 0.5 | 0.286 | 0.286 | 0.286 | 0.286 |
| | | 1 | 0.429 | 0.429 | 0.429 | 0.429 |
| | 12 | 0.1 | 0.524 | 0.524 | 0.524 | 0.524 |
| | | 0.5 | 0.571 | 0.571 | 0.571 | 0.571 |
| | | 1 | 0.619 | 0.619 | 0.619 | 0.619 |
| **0.001** | 8 | 0.1 | 0.714 | 0.714 | 0.714 | 0.714 |
| | | 0.5 | 0.714 | 0.714 | 0.714 | 0.714 |
| | | 1 | 0.190 | 0.190 | 0.190 | 0.190 |
| | **10** | 0.1 | 0.524 | 0.524 | 0.524 | 0.524 |
| | | **0.5** | **0.524** | **0.524** | **0.524** | **0.524** |
| | | 1 | 0.524 | 0.524 | 0.524 | 0.524 |
| | 12 | 0.1 | 0.667 | 0.667 | 0.667 | 0.667 |
| | | 0.5 | 0.667 | 0.667 | 0.667 | 0.667 |
| | | 1 | 0.667 | 0.667 | 0.667 | 0.667 |
| 0.01 | 8 | 0.1 | 0.476 | 0.476 | 0.476 | 0.476 |
| | | 0.5 | 0.238 | 0.238 | 0.238 | 0.238 |
| | | 1 | 0.333 | 0.333 | 0.333 | 0.333 |
| | 10 | 0.1 | 0.619 | 0.619 | 0.619 | 0.619 |
| | | 0.5 | 0.238 | 0.238 | 0.238 | 0.238 |
| | | 1 | 0.333 | 0.333 | 0.333 | 0.333 |
| | 12 | 0.1 | 0.619 | 0.619 | 0.619 | 0.619 |
| | | 0.5 | 0.762 | 0.762 | 0.762 | 0.762 |
| | | 1 | 0.333 | 0.333 | 0.333 | 0.333 |

We averaged the values of three runs for each parameter combination (i.e., repeat_1, repeat_2 and repeat_3). The results for the default parameters are shown in bold.

Table S9: Effect of parameters on Data2

| *cosdist* | *wide* | *weight* | repeat_1 | repeat_2 | repeat_3 | Average TPR |
|---|---|---|---|---|---|---|
| 0.0001 | 8 | 0.1 | 0.361 | 0.361 | 0.361 | 0.361 |
| | | 0.5 | 0.361 | 0.361 | 0.361 | 0.361 |
| | | 1 | 0.361 | 0.361 | 0.361 | 0.361 |
| | 10 | 0.1 | 0.361 | 0.361 | 0.361 | 0.361 |
| | | 0.5 | 0.361 | 0.361 | 0.361 | 0.361 |
| | | 1 | 0.361 | 0.361 | 0.361 | 0.361 |
| | 12 | 0.1 | 0.389 | 0.389 | 0.389 | 0.389 |
| | | 0.5 | 0.389 | 0.389 | 0.389 | 0.389 |
| | | 1 | 0.389 | 0.389 | 0.389 | 0.389 |
| **0.001** | 8 | 0.1 | 0.361 | 0.361 | 0.361 | 0.361 |
| | | 0.5 | 0.361 | 0.361 | 0.361 | 0.361 |
| | | 1 | 0.361 | 0.361 | 0.361 | 0.361 |
| | **10** | 0.1 | 0.361 | 0.361 | 0.361 | 0.361 |
| | | **0.5** | **0.361** | **0.361** | **0.361** | **0.361** |
| | | 1 | 0.361 | 0.361 | 0.361 | 0.361 |
| | 12 | 0.1 | 0.389 | 0.389 | 0.389 | 0.389 |
| | | 0.5 | 0.361 | 0.361 | 0.361 | 0.361 |
| | | 1 | 0.361 | 0.361 | 0.361 | 0.361 |
| 0.01 | 8 | 0.1 | 0.167 | 0.167 | 0.167 | 0.167 |
| | | 0.5 | 0.361 | 0.361 | 0.361 | 0.361 |
| | | 1 | 0.361 | 0.361 | 0.361 | 0.361 |
| | 10 | 0.1 | 0.278 | 0.278 | 0.278 | 0.278 |
| | | 0.5 | 0.361 | 0.361 | 0.361 | 0.361 |
| | | 1 | 0.361 | 0.361 | 0.361 | 0.361 |
| | 12 | 0.1 | 0.250 | 0.250 | 0.250 | 0.250 |
| | | 0.5 | 0.361 | 0.361 | 0.361 | 0.361 |
| | | 1 | 0.361 | 0.361 | 0.361 | 0.361 |

We averaged the values of three runs for each parameter combination (i.e., repeat_1, repeat_2 and repeat_3). The results for the default parameters are shown in bold.
